# Phage infection fronts trigger early sporulation and collective defense in bacterial populations

**DOI:** 10.1101/2024.05.22.595388

**Authors:** Andreea Măgălie, Anastasios Marantos, Daniel A. Schwartz, Jacopo Marchi, Jay T. Lennon, Joshua S. Weitz

## Abstract

Bacteriophage (phage) infect, lyse, and propagate within bacterial populations. However, physiological changes in bacterial cell state can protect against infection even within genetically susceptible populations. One such example is the generation of endospores by *Bacillus* and its relatives, characterized by a reversible state of reduced metabolic activity that protects cells against stressors including desiccation, energy limitation, antibiotics, and infection by phage. Here we tested how sporulation at the cellular scale impacts phage dynamics at population scales when propagating amongst *B. subtilis* in spatially structured environments. Initially, we found that plaques resulting from infection and lysis were approximately 3-fold smaller on lawns of sporulating wild-type bacteria vs. non-sporulating bacteria. Notably, plaque size was reduced due to an early termination of expanding phage plaques rather than the reduction of plaque growth speed. Microscopic imaging of the plaques revealed ‘sporulation rings’, i.e., spores enriched around plaque edges relative to phage-free regions. We developed a series of mathematical models of phage, bacteria, spore, and small molecules that recapitulate plaque dynamics and identify a putative mechanism: sporulation rings arise in response to lytic activity. In aggregate, sporulation rings inhibit phage from accessing susceptible cells even when sufficient resources are available for further infection and lysis. Together, our findings identify how dormancy can self-limit phage infections at population scales, opening new avenues to explore the entangled fates of phages and their bacterial hosts in environmental and therapeutic contexts.

## II. INTRODUCTION

Dormancy is a survival strategy found across different types of organisms, including microorganisms. Through dormancy, bacteria enter a long-term, albeit reversible state of reduced metabolic activity without cell division, enhancing survival in the face of environmental stress [1–3]. One ancient and prevalent type of dormancy in bacteria is endosporulation, which is found among *Bacillus* and *Clostridia*. Edosporulation is a complex developmental process that requires a genome duplication prior to asymmetric septum production, forespore formation, engulfment of the mother cell, which ultimately leads to the production of protein-rich endospore (or ‘spore’). Such spores are tolerant to a wide range of environmental stressors, including extreme temperatures, UV radiation, desiccation, and energy limitation [4]. Spores are abundant and have a cosmopolitan distribution (e.g., there is an excess of 10^28^ endospores in marine sediments alone [5]), can survive for extended periods of time (e.g., thousands or millions of years [6–9]), and can harbor genetic diversity given accumulation of genes in a dormant ‘seed bank’ [3, 10, 11].

Dormancy also has the potential to protect microorganisms against viral infection. Phage can account for a significant fraction of bacterial mortality [12, 13]. As a result, bacteria have evolved a wide range of intracellular and extracellular mechanisms to prevent phage infection and/or inhibit the viral replication cycle [14–17]. Well-characterized anti-viral defense systems span surface-based resistance (e.g., modifications or deletion of receptors that prevent viral infection) to intracellular defense/immunity (e.g., CRISPR/Cas or resistance-modification systems [14, 18, 19]. However, phenotypic variants of genetically identical microbes can also provide a refuge from phage infection. Examples include the decrease of phage infection of stationary phase or slow-growing bacteria [20], persister cells [21], and endospores [22]. These phenotypic obstacles are not a guarantee of protection. For example, phage T4 is capable of infecting and lysing stationary phase *E. coli* [23] while phage *λ* can infect and lyse persister *E. coli* cells even if prophage induction is inhibited [24]. In the case of *Bacillus* endospores, the protective outer layer is distinct from that of the actively growing cell and can be depleted or devoid of phage receptor binding domains. As a result, phage adsorption to spores can be reduced significantly and in some cases effectively stopped altogether [22].

The ability of dormancy to reduce adsorption of virions to cell surfaces may have consequences for population-level feedback between viruses and microbial hosts. For example, in well-mixed experimental systems, spores stabilized oscillatory host dynamics induced by phage, which reduced the extent of local population crashes [22]. The initiation and exit from dormancy may also be directly linked to interactions with viruses. For example, phage infections can trigger cell-specific dormancy initiation in *Listeria ivanovii* such that intracellular viral genomes can be eliminated during resuscitation out of dormancy [25, 26] – further mitigating the impacts of infection. Likewise, phage genomes may encode for transcription factors that change sporulation patterns in the host, potentially to circumvent host defense and lead to entrapment of viruses in spores [27–31]. Altogether, there is growing evidence dormancy initiation and revival at cellular scales is modulated as part of coevolved defense and counterdefense systems.

Scaling up the interplay of viral infection and dormancy from cellular to population scales requires accounting for feedback with the environment. Although many phage-bacteria studies are conducted *in vitro* in shaken flasks, phage infection of bacteria commonly occurs in soils, on surfaces and/or particles, and within metazoan hosts with distinct selection criteria induced by the spatial structure of the environment [32–35]. First, adjacent cells are more likely to be more closely related than if selecting cells from a population at large [36]. Second, resource environments can be patchy with variation in local availability of resources that differ substantially from the average, impacting the outcome of phage-bacteria interactions [37]. Third, lysis will not impact all other bacteria equally, instead the local propagation of phage (along with cellular debris and signals associated with infection and lysis) have the potential to influence cell fates close to sites of viral infection [18, 38–42]. All of these factors present new challenges in developing predictive models of eco-evolutionary phage-host dynamics in structured environments.

Here, we explore the interplay between dormancy and viral growth in a simplified, spatially structured environment as a means to address if and how dormancy modulates the spread of viruses within *B. subtilis* populations. To do so, we used conventional plaque assays to evaluate viral growth on a lawn of spore-forming (wild type) and non spore-forming (mutant) hosts. Phage plaques grew at the same rate initially yet produced significantly different final plaque sizes: smaller plaques in wild type vs. mutant hosts. Subsequent microscopic imaging revealed that plaques in the wild type host were surrounded by a ring of mature endospores well before the appearance of endospores far from plaque centers. As we show, this observation catalyzed the integrated development of experiments and mathematical models to explore how dormancy could form a type of collective defense that mitigates the impact of phage propagation at population scales.

## III. METHODS

### A. Experimental setup

#### 1. Bacterial strains and growth conditions

We used two 168 Δ6 *Bacillus subtilis* strains: a wild type which can sporulate, and a SPOIIE mutant in which the SPOIIE gene has been deleted (ΔSPOIIE, further referred to as the mutant strain). This mutation prevents the cell from sporulating at the asymmetrical division stage *II* which is early in the sporulation process [43] while not changing the fitness or phage infection dynamics of the WT strain [22]. We note that the Δ6 *Bacillus* strain has a reduced genome, lacks prophages, is immotile, and has reduced biofilm-forming capacity [44]. The Δ6 *Bacillus* strain is also resistant to chloramphenicol (shown as *Cm*^*R*^ in Table I).

To assist with visualization of spore development, we used a fluorescent sporulation reporter, GFP fused to a spore coat protein, under its native promoter (amyE::PcotYZ-gfp-cotZ) into both the WT and ΔSPOIIE strains. This reporter expresses relatively late in sporulation [45]. For detailed strain construction information, please consult SI Section VI A. Note that GFP is only expressed in the WT strain given that it is controlled by a sporulation-specific promoter.

Bacterial cultures were grown in Difco Sporulation Medium (DSM), supplemented with 5 *μ*g/mL chloramphenicol. DSM is a rich media that is used to obtain high sporulation rates [46, 47]. The cells were streaked from glycerol stock and grown at 37^*°*^C overnight in a shaking incubator (200 RPM) to ensure aeration of the culture. After overnight growth, cells were inoculated from a singular colony and grown under identical conditions for *≈* 5 h until they reach OD *≈* 0.5.

#### 2. Phage strains and plaque assay

We used two wild type bacteriophages of *Bacillus subtilis*: SPO1 and SPP1 (see Table I for more details on phage strains). The plaque assay protocol was adapted from the ‘tube free agar-overlay’ protocol [48]. In brief, 100 *μl* of cell culture at OD 0.5, 100*μl* of viral solution and 2.5 *ml* of 0.3% agar soft overlay at 55^*°*^C were spotted directly on the DSM 1.5% agar. Immediately after the overlay was poured, the plates were homogenized to help ensure that bacteria and viruses were evenly distributed. After the overlay sets at room temperature for 10 min, the plates were moved to a 37^*°*^*C* incubator to grow overnight.

To acquire time-lapse imaging of the plaque development, the plates were placed on top of a white LED screen in a 37^*°*^*C* room. Top-down images were captured every 5 min for 15 h. The imaging protocol was the same for end-point images in that plates were set on top of a white LED screen and a top-down image was acquired. The end-point images were taken once the plaques reached a stable state, after 15 h at 37^*°*^*C*.

#### 3. Micro plaque assay

We developed a ‘micro’ plaque assay to examine the microscale features of viral plaques. Bacterial cultures were grown to reach exponential phase as described in Section III A 1. We then prepared a viral dilution series to reach a concentration of *∼* 4 · 10^4^ pfu/ml. Concurrently, agar pads were prepared by pouring 6 mL of DSM medium with a 2% agarose concentration on a 60 × 15 mm Petri dish (based on protocol 3.3 from [49]). Using aseptic technique, the agar was cut with a scalpel into nine squares (1cm x 1 cm) (Fig. S1 panels A-D). Once the bacterial cultures reached OD *∼* 0.5, 1 mL of culture was concentrated by centrifugation at 15000 rpm for 5 min. 900 *μl* of supernatant was removed and the remaining 100*μl* before tubes were mixed in a vortex and spun down via centrifugation. The remaining 100*μl* of concentrated culture was mixed with 50*μl* of viral solution. Immediately afterwards, 2*μl* of the bacteria-virus mixture was spotted onto a 50 mm glass-bottom Petri dish (prod no 14027-20 Ted Pella). These smaller plates have a 50 mm glass bottom compatible with microscopy. A 1 cm x 1cm agar pad was placed on top of each droplet, enabling four experiments for each glass-bottom Petri dish. We then incubated the Petri dish at 37^*°*^*C* for 8-12 h before imaging (Fig. S1 panel E).

### B. Image analysis

#### 1. Time-lapse and final point image analysis

We differentiated plaques from bacterial lawn using a binarization method. Plaque centers were identified as the centroid of the largest connected component. We then iterated backwards in time to determine the centroid and plaque size in previous frames. Full details of the inward moving plaque identification algorithm with intermediate images are reported in SI Sections VI B 1 and VI B 2.

#### 2. GFP image analysis

We subtracted background fluorescence to compute the average GFP intensity relative to the center of the plaque. The corrected image was used to identify the center of the plaque. Finally, the distance between each pixel and the center was calculated to obtain panel B in Fig. 3; full description provided in SI section VI B 4.

**FIG. 1:**
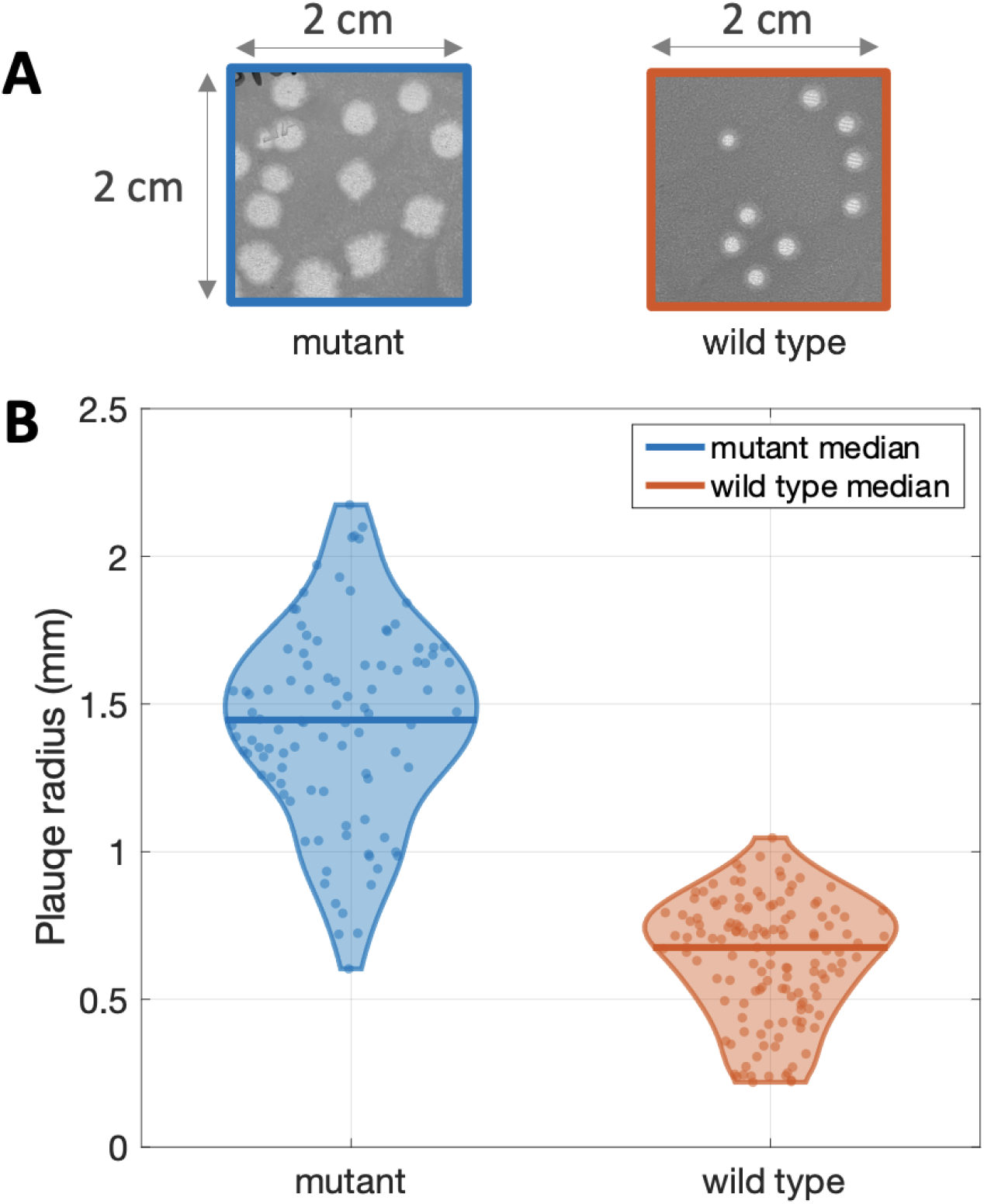
Plaque sizes of phage SPO1 with mutant and wild type host. (A) Square cropped sections of size 2cm x 2cm from mutant (left) and wild type (right) plaque assays are shown. Both plaque assays were carried as described in section III A 2. mutant and wild type plaque assays where prepared in parallel and have the same initial and growth conditions. (B) Final plaque size of the two plaque assays shown in panel A. The images were analyzed using binarization and watershed algorithm (see SI section VI B 1). The plaque sizes (98 plaques for mutant and 137 for wild type) were plotted using a violin plot and individual points are shown as a scatter plot. The median mutant and wild type plaque sizes are shown through the horizontal bold lines. Mutant plaques have a radius of 1.43 *±* 0.33*mm* and wild type plaques have a radius of 0.64 *±* 0.2*mm*. A two sample t-test was performed to reject the hypothesis that the two distributions have equal means with a p-value less than 10^*−*3^.

**FIG. 2:**
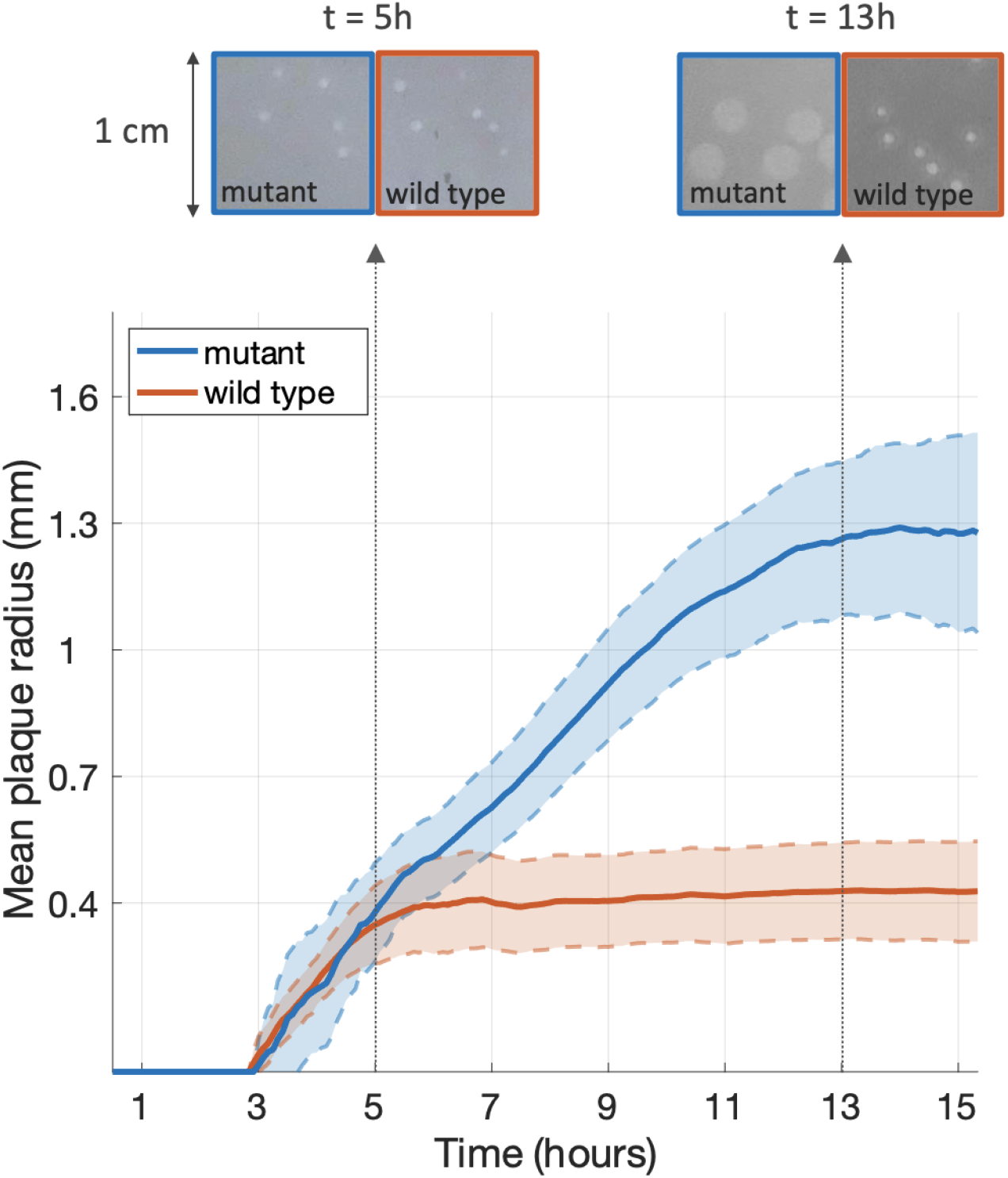
Time lapse of plaque growth for mutant and wild type host. A traditional plaque assay was performed at 37^*°*^ C and images were captured every 5 minutes over a period of 15 hours. The plaque sizes were estimated for each time point (46 plaques for mutant and 100 plaques for wild type) using image analysis techniques summarized in Section III B and described in detail in SI section VI B 2. Cropped sections from the time-lapse are shown at 5 and 13 hours for parts of the mutant and wild type plates. The mean over all plaques is shown in the solid line and the standard deviation is shown in the shaded area. Intermediate images of the independent plaque trajectories are shown in Figure S5.

**FIG. 3:**
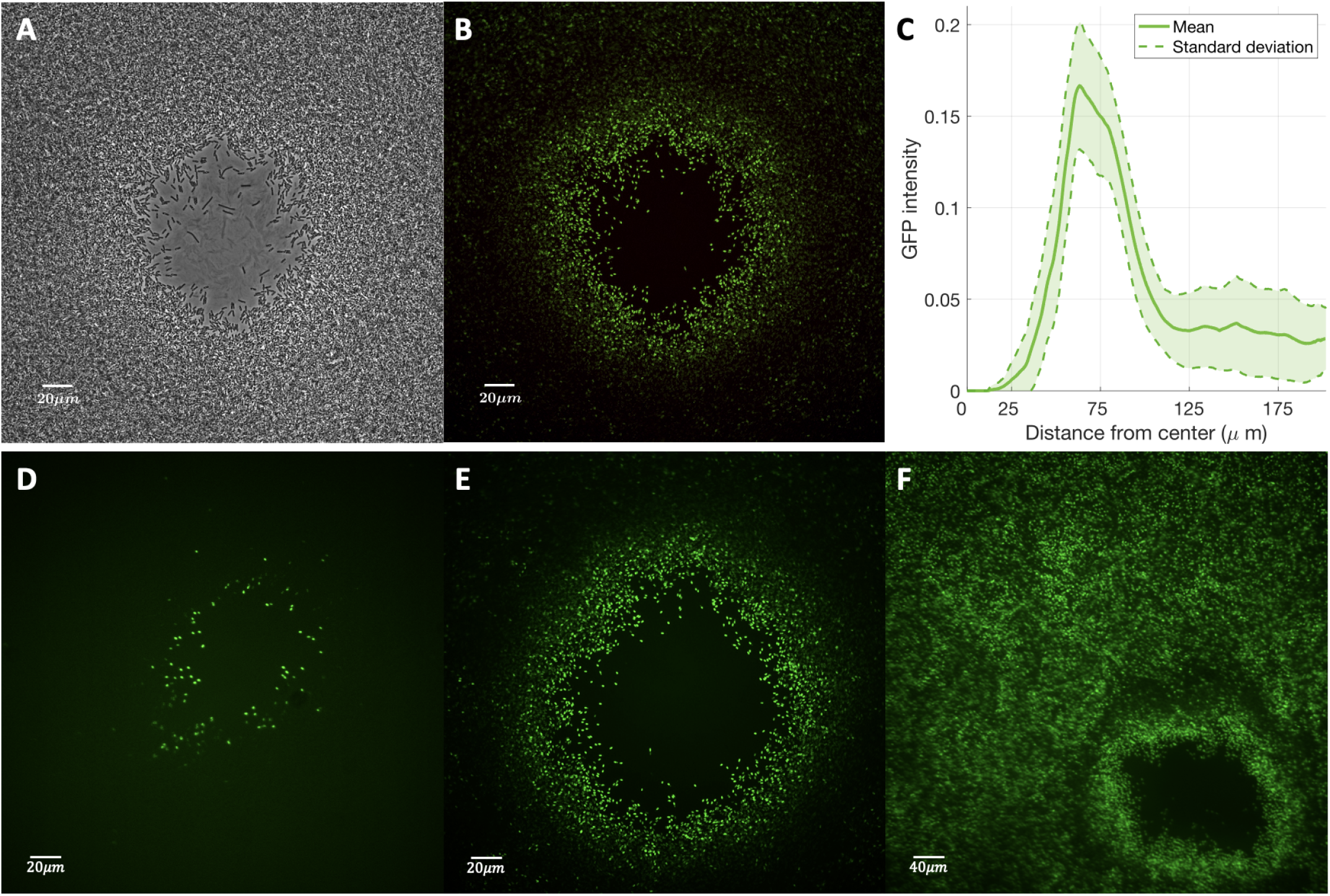
Sporulation is enhanced around plaque edges. Bright-field image of micro plaque assay is shown in panel A and fluorescent images are shown in panels B, D, E and F. Images A, B, D and E were taken at 40x magnification and image F at 20x magnification. The micro plaque assay was carried out with phage SPO1. The GFP image was adjusted to remove background fluorescence and enhance contrast (see section VI B 4). (C) GFP analysis of image in panel B. An averaging filter is applied on the image in panel B to obtain an adjusted GFP image. The green component in every pixel is then computed based on the distance to the center of the plaque (see SI section VI B 4). Distance to the center of the plaque is shown on the x-axis in *μm* and GFP intensity in the adjusted image is shown on the y axis. The solid line represents the mean and the shaded area is the standard deviation. Panels D, E and F show micro plaque assays at different time points from earliest in panel D to latest in panel F. The image in panel D was taken within the first 8 hours when only cells in the vicinity of the plaque were sporulating. The image in panel F was taken after 12 hours when most of the cells have transitioned to spores.

### C. Mathematical models and simulations

#### 1. Mathematical modeling

We developed a series of nonlinear partial differential equation models of phage-bacteria interactions in a spatially explicit context, building upon mechanisms described in well-mixed [50] and structured [41] populations. In this model, susceptible bacteria *S* grow on resources *R* and can be infected by viruses *V* yielding infected cells *I*. Infected cells can lyse releasing new viruses. Susceptible cells can also transition to dormant spores *D* – phage cannot adsorb to spores [22]. To account for different modes of dormancy initiation, we developed three different model variants: Model R, Model V and Model M (Fig. 4) that differentiate the mechanism underlying dormancy initiation: R - depletion of resources; V - interaction with virus particles and, separately, resource depletion; M - interaction with lysis-associated molecules and, separately, resource depletion. Initially, susceptible cells and resources are uniformly distributed while viruses are concentrated at the central point. There are no dormant or infected cells at the beginning of a simulation. In all model variants, we explicitly account for the spatiotemporal dynamics of populations.

**FIG. 4:**
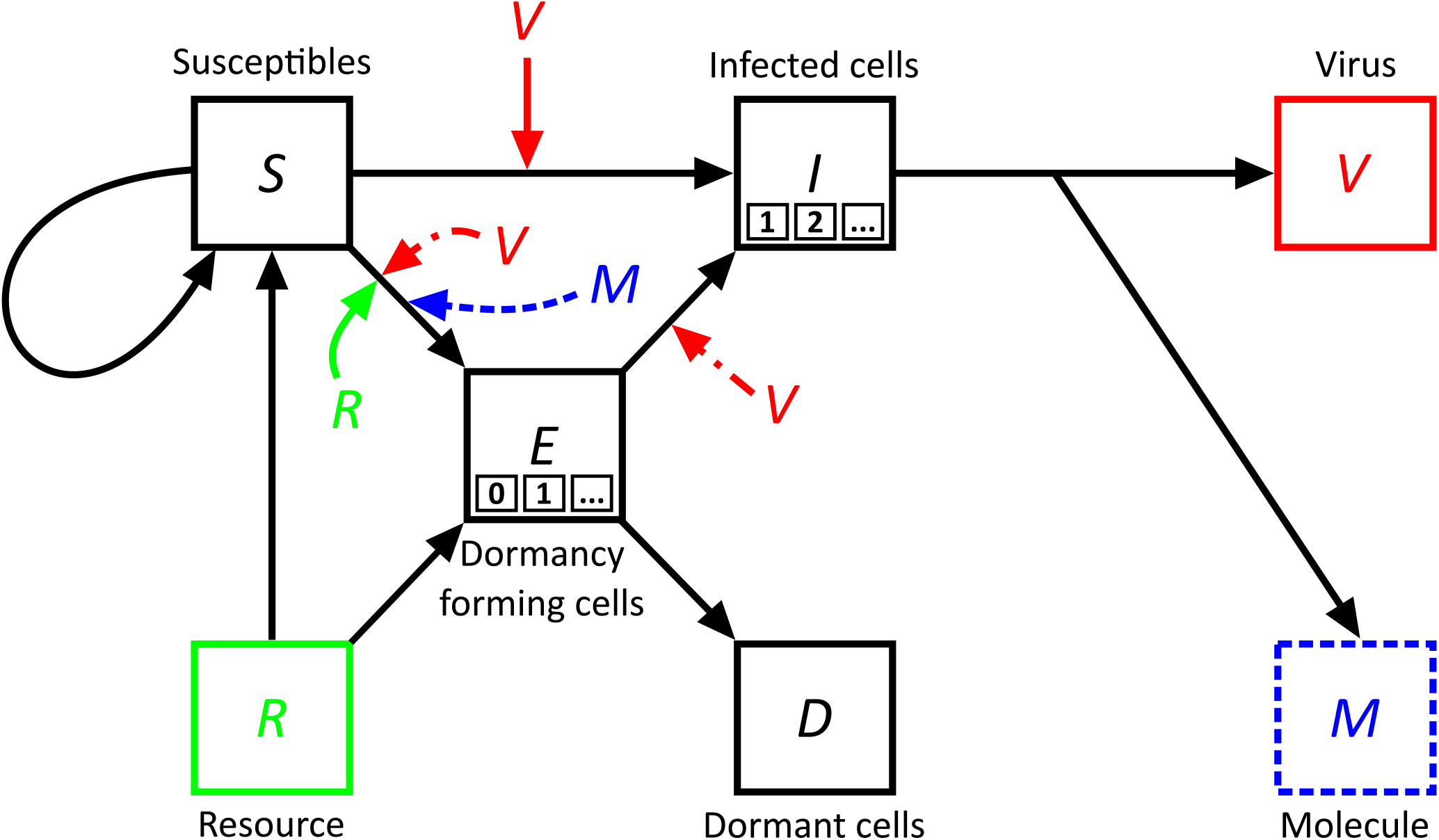
Schematic comparison of Models R, V, and M. Each shape represents a specific element in the system. Squares correspond to explicit limited resources (R), susceptible bacteria (S), infected bacteria (I), dormant bacteria (D), bacteria transitioning to dormancy while exposed to viruses (E), free viruses (V) and signaling molecules (M). In the E and I boxes, the smaller boxes signify the potential number of E and I states, denoted as *n*_*E*_ and *n*_*I*_ respectively. The number of states defines the duration that cells spend in that state (see Methods-Mathematical Modeling III C 1). For the E states, *n*_*E*_ can be set equal to zero, which corresponds to a direct transition from the susceptible state to the dormant state. The arrows indicate interactions, with their direction showing causal effects. The central elements of Model R, Model V, and Model M are highlighted in green, red, and blue respectively. These are, the Resource for Model R, where we assume resource-only dependent dormancy, the Virus for Model V, where dormancy can be triggered by viral contact aside from starvation and the Molecule for Model M, where we assume an infection-associated molecule is an additional, potential initiator for dormancy. Elements with solid lines are shared across all models, whereas elements with dashed lines are unique to specific models and correspond to the model of the same color.

#### Model R - Resource-only dependent dormancy

Model R assumes that sporulation in *Bacillus subtillis* is triggered by starvation [3, 43, 51], specifically, dormancy is initiated at a low rate when resources are high and at a much higher rate when resources are depleted [52, 53]. The associated partial differential equations are as follows:

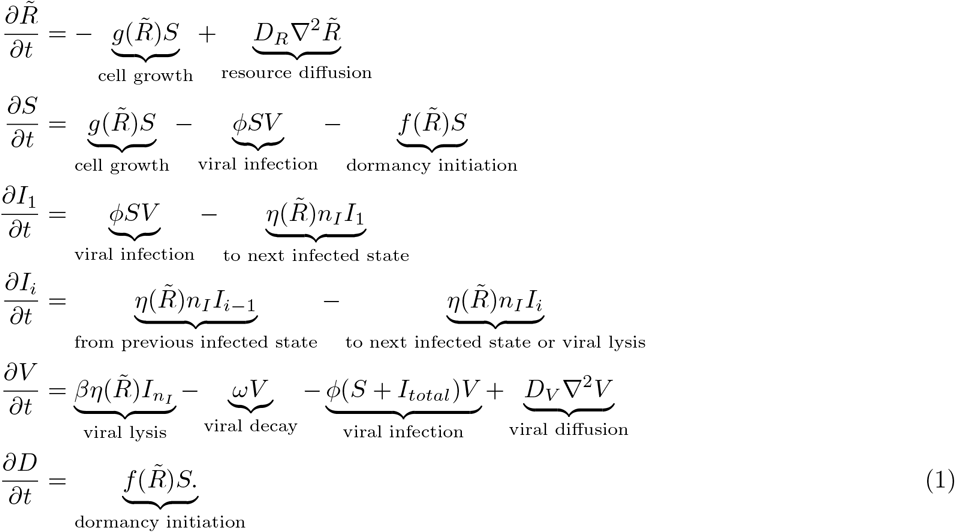

Where 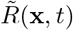 is the rescaled density of resources, expressed in units of cells/mL. The rescaling is performed as 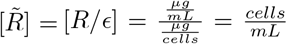, where *ϵ* denotes the rate of resource to bacteria conversion measured in *μ*g/cells (see Table II and Parameter estimations for details). Similarly, *S*(**x**, *t*), *I*(**x**, *t*) and *D*(**x**, *t*) are the densities of susceptible, infected, and dormant bacteria (i.e., spores), respectively, each in units of cells/mL. *V* designates the density of phage in viruses/mL. Here, immotile bacteria grow by consuming resources while resources and viruses diffuse at distinct rates. Susceptible cells can be infected by virulent phage with infection rate *ϕ* or become dormant if the density of the resources is sufficiently low. Once infected, the cells go through a series of sequential stages *n*_*I*_ of infected states *I*_*i*_ (the sum of which is given by 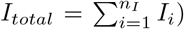, resulting in an effective latent time delay which follows an Erlang distribution. After the latency period, the infected cells burst yielding new free viruses with burst size *β*. Virus particles can adsorb to cells at a rate *ϕ* and decay with rate *ω*. The bacterial growth rate 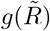, dormancy rate 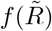 and latent time 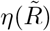 are dependent on available resources and are given by:

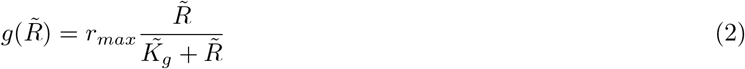

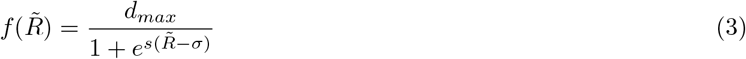

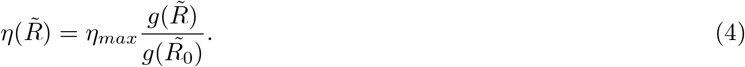

The growth rate function follows the Monod equation where *r*_*max*_ is the maximum bacterial growth rate and 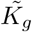 the rescaled Monod constant (similarly to R, 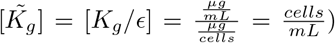. The dormancy rate function is assumed to be solely dependent on resources but with sporulation starting effectively only when the resource density is close to a fraction *σ* of the Monod constant (for Model R 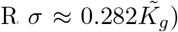. The ‘sharpness’ of the transition to dormancy is determined by the value of parameter *s*. Finally, we choose a resource dependent lysis rate that is proportional to the growth rate *g*(*R*) [54, 55]. This choice is justified by the fact that the plaques stop growing as bacteria have exhausted the available resources. Full details of parameter estimation are in Section VI C 2 and model parameters are in Table II.

#### Model V - Contact mediated dormancy

In Model V, dormancy can be initiated via starvation or with a probability *p ∈* (0, 1) upon interacting with a virus particle. This yields a family of (*p, σ*) combinations that exhibit identical plaque growth dynamics, with the (*p* = 0, 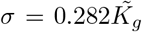) case corresponding to Model R. In order to account for a time delay for spore formation we introduce a series of multiple sequential stages *n*_*E*_ to become dormant [53]. Hence, the dormancy initiation time follows an Erlang distribution. The transition to dormancy is immediate for *n*_*E*_ = 0, while a delay is introduced for 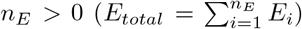. The net transition rate between the E-states is *λn*_*E*_, where 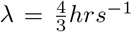, so that the mean transition rate remains constant with varying number of states. In this model formulation, cells transitioning to dormancy can become infected by phage, whereas spores cannot be infected (see Discussion for elaboration on the consequences of relaxing this assumption). All other parameter values are retained across models R and V (as shown in Fig. 4 and Table II. The new terms in Model V are listed in red text, the remainder are shared with Model R:

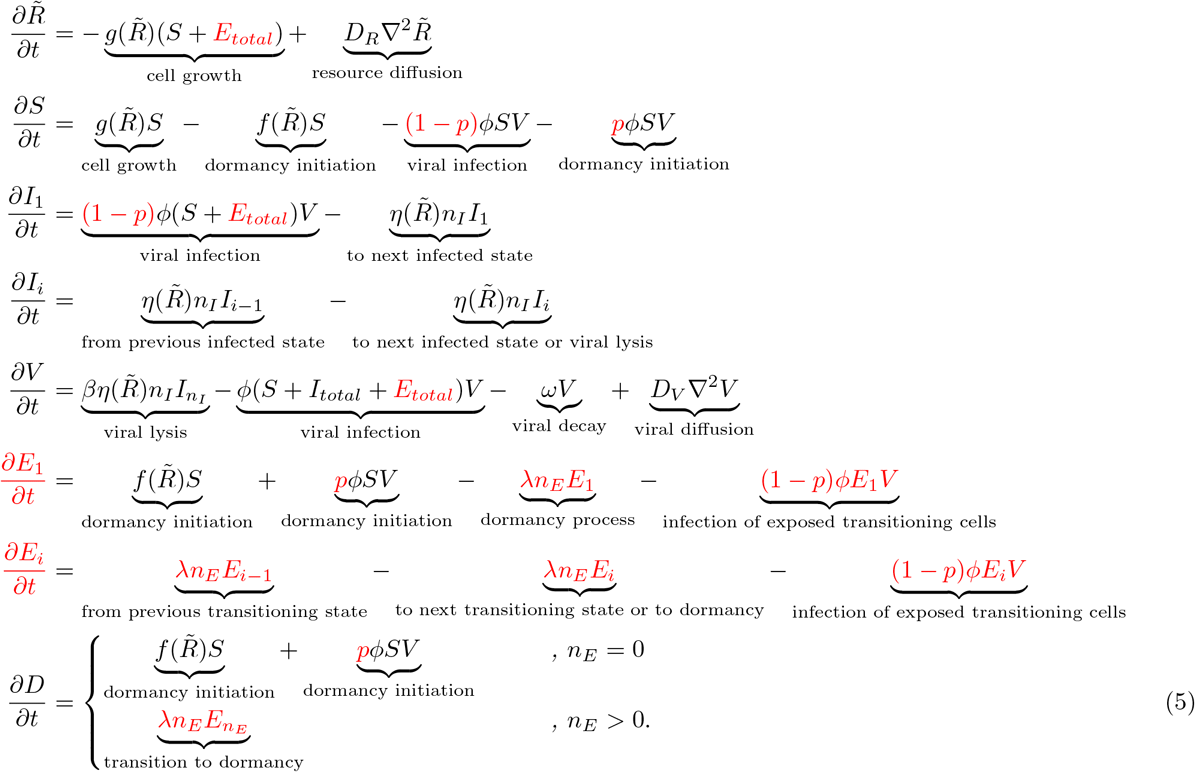

#### Model M - Dormancy mediated by lysate-associated molecules

Model M represents a scenario where sporulation can be triggered by diffusible molecules released upon cell lysis. Additional parameters include the rate at which these lysate-associated molecules trigger a susceptible cell to become dormant, *μ* = 5 · 10^*−*11^mL*/*(hrs · (cells)), the number of molecules released upon lysis *m* = 10^4^ molecules/cell, and the diffusion constant *D*_*M*_ = 4 · 10^5^*μ*m^2^/hrs. The parameters 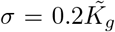 and molecular concentrations are in cells/mL - representing the equivalent density of cell-to-spore transitions that molecules can trigger. The following system of equations expands Model R and Model V (see equations sets (1) and (5) for comparison with the black and red elements in Model M respectively) with the blue terms that are introduced in Model M.

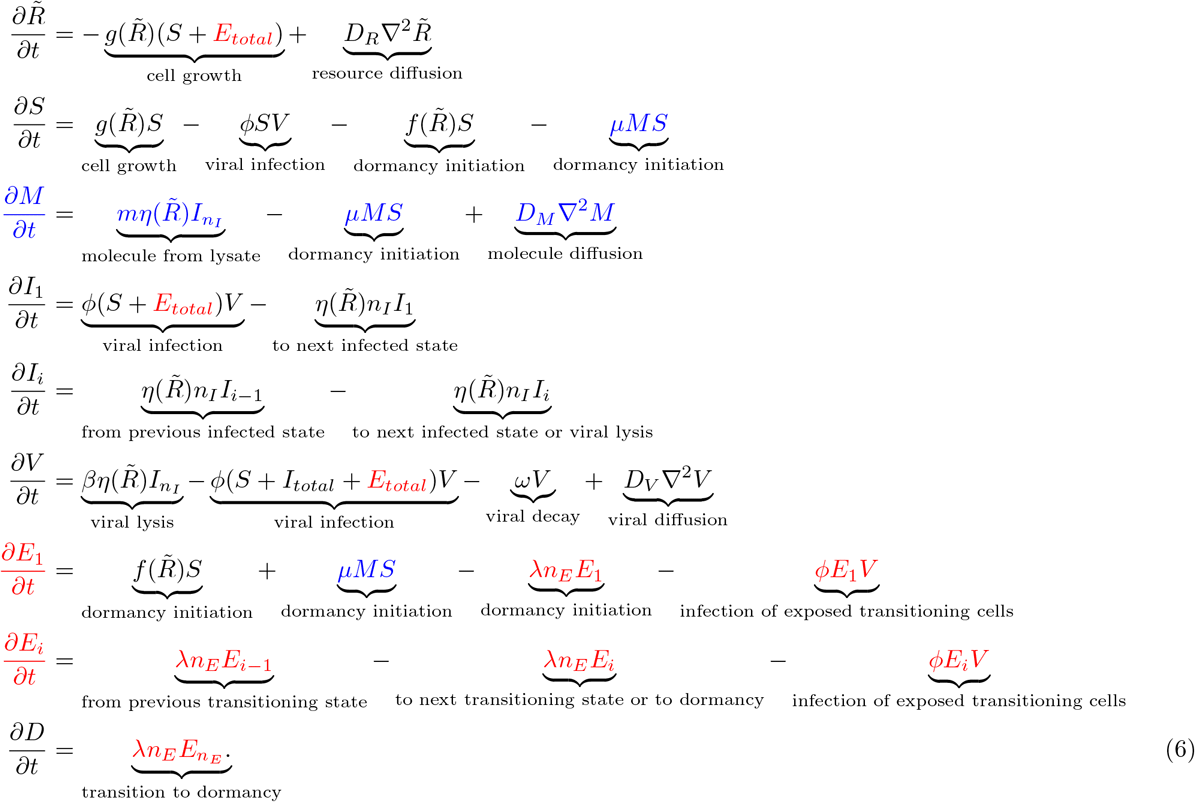

### D. Data and code availability

All simulations were carried out in Python (Jupyter notebooks) and image analysis was carried out in MATLAB v 2020a. Scripts and data are available on Github at https://github.com/WeitzGroup/Plaqueearlysporulation.

## IV. RESULTS

### A. Plaque growth dynamics in sporulating vs. non-sporulating hosts

We performed standard plaque assays of phage SPO1 with wild type and mutant hosts at 37^*°*^ C over a 15 h period, as described in section III A 2. In both cases, phages proliferated and generated macroscopic zones of clearance, i.e., plaques. On average, the radius of mutant plaques was *∼* 2.2 fold greater than the radius of wild type plaques (Fig. 1 A). Mutant plaques radii were 1.43*mm ±* 0.33*mm* and wild type plaques were 0.64*mm ±* 0.2*mm* (t-test, *p <* 10^*−*3^). We assessed the generality of this finding by comparing plaque sizes on sporulating vs. non-sporulating hosts using a different bacteriophage, SPP1. In this case, we observed a 3.5 fold plaque radius reduction for bacteriophage SPP1 *p <* 10^*−*3^, given mutant plaque radii of 1.22*mm ±* 0.15*mm* and wild type plaque radii of 0.35*mm ±* 0.12*mm* as shown in Fig. S2). Hence, we observe a similar phenomena across multiple phage types: plaque size is reduced by 2-4 fold in a population of sporulating bacteria vs. non-sporulating bacteria.

In order to further quantify plaque growth dynamics on sporulating vs. non-sporulating hosts, we recorded a time-lapse of the SPO1 plaques. We extracted the plaque sizes at every 5 min over 15 h (see Methods section III B and SI section VI B 2), Fig. 2). Both mutant and wild type plaques started growing at the 3 h mark and continued to grow for 2 h. However, the wild type plaques reached a plateau shortly after 5 h, whereas the mutant plaques continued to grow until reaching a plateau at 13 h (see differences in wild type and mutant dynamics, Figure 2). The difference in plaque growth duration is associated with differences in plaque sizes – mutant mean plaque size of 1.28 *±* 0.24*mm* vs. wild type mean plaque size of 0.43 *±* 0.12*mm*. Notably, the reduction of plaque size in sporulating hosts across different phage (Figures 1 and 2) are robust to experimental conditions required for both time-lapse and endpoint measurements.

We distinguish three phases of plaque development: a lag phase in which plaques are not visible, an enlargement phase in which plaques appear to be growing at a constant rate and a termination phase in which plaques stop growing, presumably due to a halt in phage replication [38]. We fit a linear function to the enlargement phase for each mutant and wild type trajectory to obtain 46 growth rates for mutant plaques and 100 growth rates for wild type plaques. The mutant mean growth rate is 117 *±* 26*μm/hr* and the mean wild type growth rate was 136 *±* 69*μm/hr*. We conducted a bootstrap resampling of growth rates and find that a fraction *p* = 0.066 of 10^5^ slope differences in randomized groups exceed that of the observed difference of 19 *μ*m/hr (see Fig. S6). Hence, the growth rate differences of the two bacterial strains are not significantly different, despite the observation that the final mean plaque size is almost three times larger for mutant compared to wild type. The early cessation of plaque growth in sporulating vs. non-sporulating hosts suggests that there are additional mechanisms limiting the ability of phage to lyse hosts.

### B. Sporulation analysis across plaque transects

In order to explore the basis for our observation of early cessation of plaques in sporulating hosts, we developed a microscopic plaque assay to quantify the levels and locations of mature endospores relative to plaque centers. Fig. 3) contrasts the bright-field image with the resulting microscopic plaque assay (see section III A 3). The bright-field image of a plaque is shown in panel A; it resembles a traditional viral plaque with a clearing in the middle surrounded by a bacterial lawn. However, as shown in Fig. 3B, there is an absence of spores near the plaque center and an enhancement of spores around the edge of the plaque compared to the rest of the bacterial lawn. This enhancement of spore density near plaque edges relative to the rest of the bacterial lawn was captured 8 h after the initiation of the experiment and represents a critical transient that would otherwise be missed in end-point analysis. Indeed, after a prolonged time period of *∼* 16*h*, the bacterial lawn reached sporulation levels similar to those found around the plaque edge (Fig. 3 F). This suggests that sporulation is triggered earlier in cells that are close to regions with enhanced viral-induced lysis. We note that the enhancement of sporulation around plaque edges is robust, and observed across multiple plaques in multiple experiments. Fig. 3D-F shows examples of multiple stages of the enhancement of spores around plaque edges, including at later stages where the densities of cells around plaque edges matches that of intensities in the background (Fig. S7).

The distribution of spores relative to plaque centers was further quantified by measuring the GFP intensity as a function of the distance to the center of the plaque (Fig. 3C). In each case, we identified plaque centers and then averaged GFP intensity in an annulus of a fixed distance from the center. The lowest GFP level was obtained when the distance to plaque center was less than *∼* 25*μm*. GFP expression increased between *∼* 15*μ*m-100*μm* before decreasing to background levels far from plaque centers. This quantitative enhancement of GFP intensity associated with the early emergence of spores is localized to plaque centers – again emphasizing the importance of a dynamical feedback between a spreading viral population associated with localized lysis and the early onset of endosporulation. These findings also raise the question on what cellular mechanisms are compatible with the emergence of a collective ring of spores outside plaque edges associated with early plaque termination.

### C. Modeling plaque growth given dormancy initiation by resource depletion

We developed and analyzed a mathematical model of phage spreading across a bacterial lawn including either sporulating (*S*^+^) or non-sporulating (*S*^*−*^) bacteria to better understand what factors affect plaque growth. The mathematical model initially included bacteria growth, viral infection and lysis, the potential diffusion of virus particles and resources as well as the initiation of cellular dormancy due to resource depletion (full details in Sec. III C 1). Our objective was to compare simulated plaque spreading dynamics with observations arising from both macroscopic and microscopic plaque assay experiments. As noted, macroscopic plaque analysis revealed that plaques are substantially smaller when grown on sporulating vs. non-sporulating hosts due to an early termination of plaque growth rather than a change in plaque growth speed. Further, microscopic plaque analysis revealed the early emergence of a ring of endospores around the edge of plaques before that found in the bulk. In developing this partial differential equation (PDE) model, we first evaluated the scenario where starvation induces dormancy. We term this the resource-dependent model or Model R, such that sporulation is enhanced at low resource concentrations and suppressed at high resource concentrations [3, 43, 51].

The PDE model of phage-bacteria interactions in a depleting resource environment generates a spreading wave. Model R reproduces the macroscale observation of reduced plaque size in sporulation vs. non-sporulating hosts (contrast Fig. 1, experimental results with Fig. 5A, simulation results). The underlying reason is that viruses spreading in the wildtype population have fewer available cells to infect compared to those in the mutant population. Furthermore, Model R also reproduces the early, equivalent plaque growth dynamics and the early cessation of expansion observed in plaques associated with the *S*^+^ vs. *S*^*−*^strain, as corroborated by Fig. 5B. In a model with starvation-induced sporulation, dormancy typically starts from the exterior part of the plate where bacteria growth is not controlled by phage. Therefore outside the plaque, resources are abruptly depleted sooner, triggering dormancy in a large fraction of cells uniformly distributed out of the reach of phage activity (i.e., outside the plaque). Phage originating from the plaque’s center will eventually reach a region with dormant cells present at relatively higher densities, which prevents further infection and amplification of plaque spread leading to smaller plaques for the *S*^+^ strain.

**FIG. 5:**
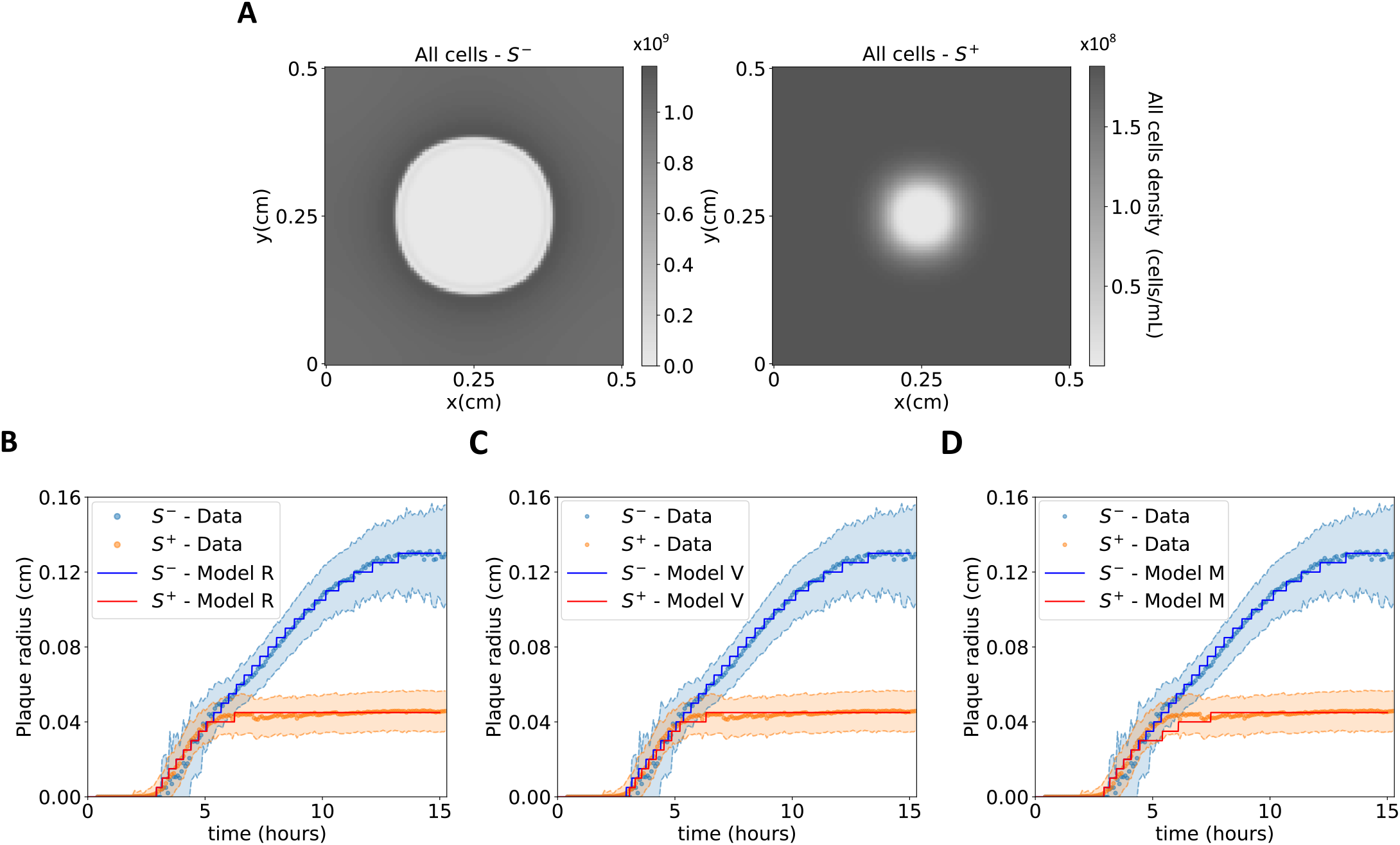
Impact of sporulation on plaque size for Models R, V, and M. (A) A snapshot from the spatial simulation of bacterial densities in Model R, taken at 15 hours, corresponding to the terminal point of the time-series shown in panel B, using parameters from Table II. (B-D) Overlay of experimental data and computational simulations shows the time evolution of the mean plaque radius for the *S*^*−*^ and *S*^+^ strains in Models R, V, and M, respectively. In the simulations, a plaque is defined as the area where cell density is less than 10% of the maximum density at that time point. The results exhibit robustness to variations in that threshold, attributed to the abrupt density changes at plaque boundaries. In panels C and D, simulations for Model V and Model M respectively are performed with *n*_*E*_ = 10. Results are replicable across different values of *n*_*E*_ (see Fig. S9 for the *n*_*E*_ = 0 case).

Despite the agreement with macroscale observations of plaque dynamics, Model R incorrectly predicts that sporulation density should continue to increase away from plaque centers into the bulk bacterial lawn in contrast to micro plaque assays (see Fig.6A,B). This increase arises because resources are consumed equally in the absence of viruses triggering a uniformly distributed background of dormant cells. With increasing proximity to the plaque there are fewer susceptible bacteria due to lysis, leading to higher resource availability and lower dormant cell densities. At the edge of the plaque there is a precipitous decrease in bacteria density as all cells are killed, corresponding to a sharp decrease in dormant cells (see Fig. 6A,B). Therefore, in this model the decrease in resources is fastest farther from regions of viral infection and lysis, i.e., from the ‘outside’ of the plate towards the phage-occupied center. Model R’s prediction that spore densities should increase with distance from the plaque center both transiently and at the end point suggests that processes beyond starvation are inducing spore formation in the experimental system.

**FIG. 6:**
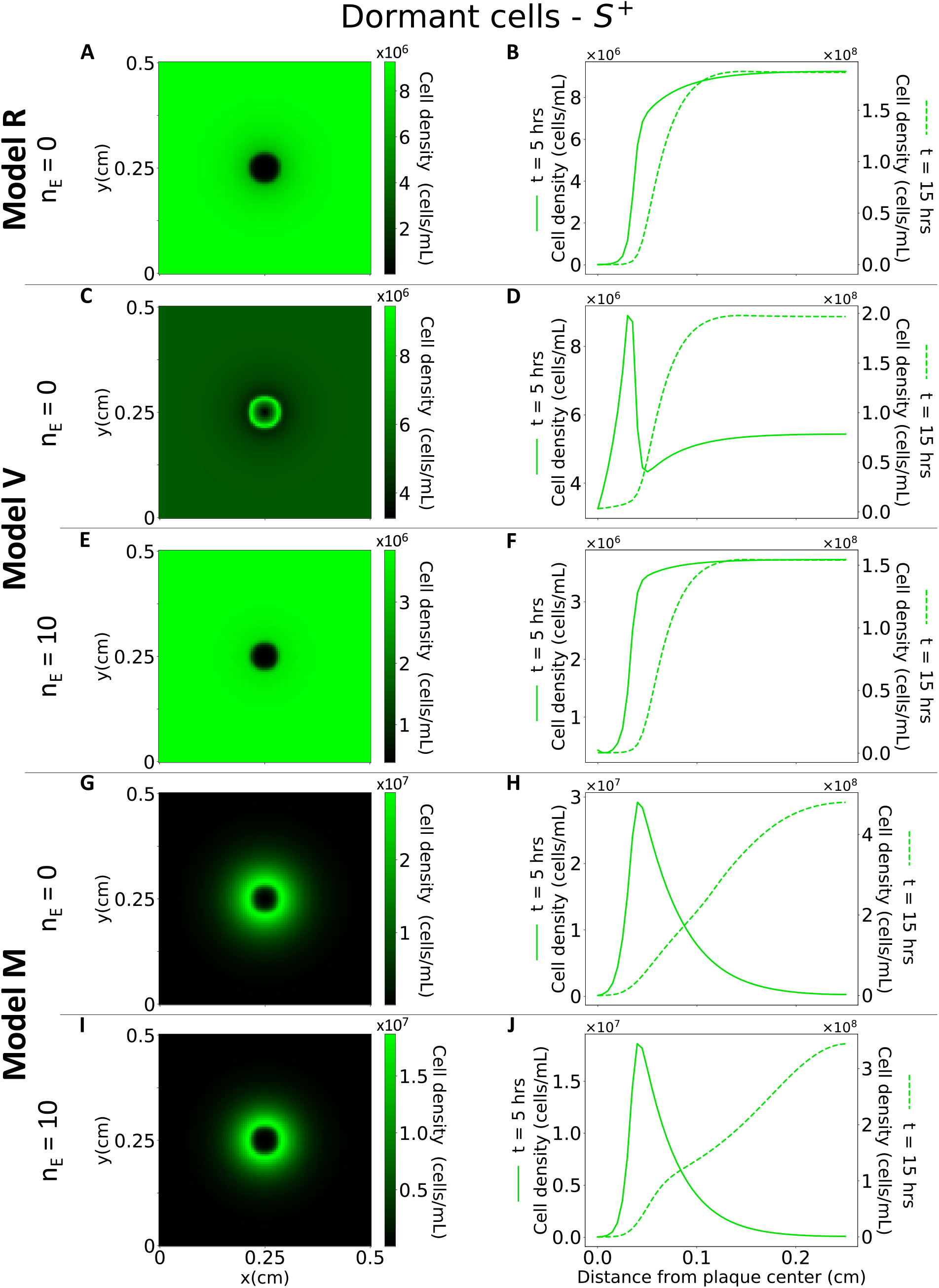
Simulated plaque growth and distribution of dormant cells for the *S*^+^ host at t = 5 hrs after phage addition. The arrangement of the simulation results aligns with that presented in Fig. 3B,C. Rows are organized by model type as results for Models R, V and M. The left column (panels A, C, E, G, I) depict dormant cell densities’ 2-D spatial distribution. The results correspond to t = 5 hrs, which is centered in the time frame of t *≈* 3 - 7 hrs during which the transient peak in sporulation around the plaque emerges in our simulations. The right column (panels B, D, F, H, J) show the distance profile of dormant cells distribution starting from the center of the plaque. The solid line (primary y-axis) corresponds to results at time = 5 hrs and is effectively a 1-D slice of the left panel at the same row, showing the radial distribution of dormant cells starting from the center of the plaque and moving outwards. The dashed line, plotted on the secondary y-axis, represents the progression of the solid line at 15 hours.

### D. Modeling phage plaque growth given direct, infection-triggered sporulation

Next, we evaluated Model V in which bacteria dormancy is triggered by two distinct mechanisms: (i) resource depletion, as in Model R; (ii) infection by phage. Like Model R, Model V can reproduce quantitative plaque growth dynamics, insofar as plaques grow at the same speed on both sporulating and non-sporulating hosts yet stop earlier on sporulating hosts vs. non-sporulating hosts (see Fig. 5C and Fig. S9). In Model V, viruses diffusing at the plaque front can trigger cellular dormancy amongst newly infected cells before population growth-induced depletion of resources. Hence, viruses spreading on sporulating hosts will be diluted out through encounter with hosts, eventually leading to the early cessation of plaques relative to that of viruses growing on mutant hosts that cannot form endospores.

Although Model V and Model R may both be compatible with the macroscopic plaque growth dynamics, they differ in the emergence spatiotemporal profile of spores. In Model V, the spore density increases outwards from the center as there are not enough cells inside the plaque for appreciable infections to take place. Likewise, further away from the plaque, there is less phage-induced dormancy. If resources have not dropped yet below the critical value of *σ*, the spore density drops, reproducing the experimentally observed peak at the edge of the plaque (see Fig. 6C,D). Similarly to our experiments (Fig. 3D-F), the peak in our simulations is transient, emerging at *≈* 3 h and persisting until t *≈* 7 h. Given that there is a 4-5 h interval between the transition to dormancy and the expression of GFP, this simulated onset at t *≈* 3 h closely aligns with our experimental observations, which show the peak at t *≈* 8 h.

The localized sporulation peak around plaque edges is transient given that dormancy is initiated in ‘far-field’ cells distant frmo plaque centers as a result of resource depletion. Consequently, Model V will relax to the same steadystate spore density profile as in Model R. Therefore Model V can qualitatively reproduce the transient sporulation enhancement around the plaques as manifested in the comparison between the experimental (Fig. 3) and the modeling results (Fig. 6C,D)) at t = 5 h. However, Model V also produces dormant cells from the early stages of the plaque formation, with spores within the plaque building up right away from its center (see Fig. 6C,D). Conversely, in the experiments, the central region of the plaque typically does not have spores (see Fig. 3).

Thus far the modeling results in Fig. 6C,D assume direct conversion from susceptible cell (S) to dormant cell (D). We next evaluated the robustness of our findings for Model V by including a more realistic delay before the onset of dormancy, through the introduction of a sequence of multiple intermittent states *n*_*E*_ (see Methods, Model V). Given the inclusion of a delay before dormant cells resist subsequent viral infection [53], we find that Model V continues to reproduce plaque growth dynamics (see Fig. 5C and Fig. S9A) but does not generate a ring of increased dormant cells around the plaque (see Fig. 6E,F and Fig. S10). In Model V, bacterial initiation of dormancy can be the result of direct interaction with phage. Hence, cells are not preconditioned to initiate dormancy required to generate an annulus of dormant cells. In aggregate, once an explicit latent period is introduced then although virus spread may decrease, the net result is – like Model R – that resource scarcity becomes the primary driver of dormancy and the spatiotemporal profiles of spores are incompatible with experiments.

### E. Modeling phage plaque growth given infection-associated molecule-triggered sporulation

An alternative hypothesis is that dormancy may be initiated inside cells that have not yet been infected by phages due to the uptake of host metabolites, host lysis signals, and/or viral proteins that diffuse through the environment faster than virions. Hence, we evaluate Model M in which dormancy can be initiated through one of two mechanisms: (i) resource depletion, as in Model R; (ii) uptake of infection-associated molecular triggers. Model M can reproduce the observed reduction of the plaque size in systems with sporulating vs. non-sporulating cells as well as quantitative plaque growth dynamics, as shown in Fig. 5D and Fig. S9B. Notably, Model M also successfully replicates the transient peak of sporulation around the plaques for the same time-period as Model V with *n*_*E*_ = 0 and with a delayed transition to dormancy (unlike Model V; see Fig. 6G-J). Central to this observation is the assumption that the signaling molecule diffuses faster than viruses. At a certain distance from the center, cells encountering the molecule initiate and complete differentiation into mature endospores resistant to infection at densities high enough to stop plaque propagation. Towards the center, lysis kills cells that are transitioning to spores, reducing their density. Farther out than the peak, the spore density decreases again given that the diffusing molecule associated with sporulation initiation will not reach sufficiently high densities yet. Unlike either Model R or Model V, Model M can also reproduce a central region of the plaque, empty of spores and cells (see Fig. 6G-J) as observed experimentally (see Fig. 3).

## V. DISCUSSION

In this study, we tested the effect of sporulation on phage plaque formation within *B. subtilis* populations. Macroscopic plaque analysis revealed that phage plaques grow at indistinguishable rates in sporulating (wild type) vs. non-sporulating (mutant) strains. However, plaques grown on wild type populations exhibit rapid and earlier cessation of growth compared to those grown on the mutant strain, leading to 2-3 fold smaller plaque sizes. Via a microscopic imaging assay of the interior and boundary of plaques, we found that sporulation is enhanced around plaque edges well before resource depletion leads to the emergence of spores in the population as a whole. Together our findings suggest that the emergence of enhanced sporulation in proximity to lysis events serves as a collective, protective mechanism against further phage dispersion and lysis.

We were able to recapitulate the spatiotemporal dynamics of plaque front growth and the emergence of an annulus of spores around plaques with a spatially explicit model of virus-microbe interactions that incorporates lysis-associated initiation of dormancy. Notably, models in which dormancy is initiated due to resource depletion and/or infection alone do not recapitulate the formation of an annulus of spores around plaques associated with early plaque termination. Instead, the emergence of an annulus of spores surrounding phage plaques as observed in micro plaque assays can arise given relatively faster rates of the diffusion of infection-associated molecules that can trigger dormancy initiation compared to virus particle diffusion (see Figure 7). The rationale is that faster diffusion of molecules enables cells to initiate dormancy in advance of virion arrival and well before resources are depleted far from plaque centers. Hence, the combined evidence from plaque assays, microscopic imaging, and mathematical models shows that dormancy is locally enhanced by lysis events and that this enhanced dormancy can, in turn, slow the propagation of viral populations.

**FIG. 7:**
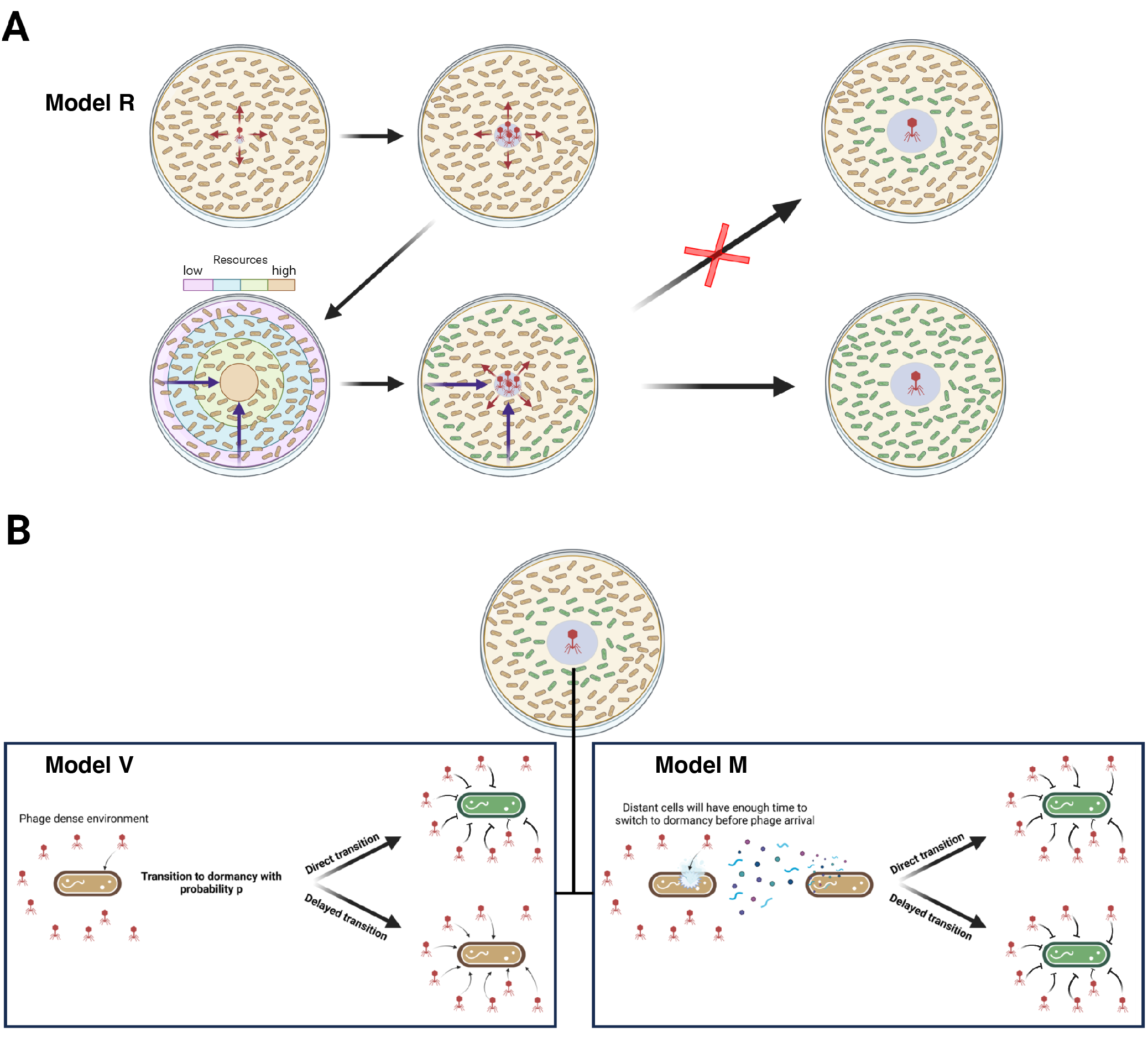
Schematic of lysate-associated termination of plaque fronts and the emergence of sporulation annuli. (A) In classic resource-limited models of dormancy initiation, resources should be depleted more rapidly in regions without phage clearance, as cells grow to higher density. Hence sporulation should theoretically proceed from the ‘outside’ inwards rather than leading to an annulus as observed experimentally. (B) Alternative sporulation-induced mechanisms include direct initiation by infection and indirect initiation through a signalling molecule. If virus contact initiations dormancy, then it is possible for a sporulation annulus to emerge insofar as the transition is immediate (i.e., cell takeover fails). However, given that protective benefits of dormancy typically accrue after a maturation delay, then contact alone is insufficient to yield a sporulation annulus. In contrast, the earlier arrival of a signaling molecule in the viral lysate (that diffuses faster than phage particles) would catalyze earlier dormancy initiation and generate a sporulation annulus around plaques whether dormancy initiation was direct or delayed. *Note: in all panels, brown denotes active cells and green denotes mature spores*.

We posit that there are multiple classes of cellular mechanisms that could give rise to enhanced sporulation around plaques. First, it has recently been shown that a specific type of siderophore (coelichelin) released by a competitor *Streptomyces* strain can reduce iron availability within *B. subtilis* populations, which in turn limits sporulation [56]. While this interspecies effect alters the outcome of competition, iron availability may also influence spore initiation with implication for phage infection. For example, viral lysis within a developing plaque may increase the local concentration of bioavailable iron which is then used by *B. subtilis* to make endospores — leading to a strongly, self-limiting plaque. Second, in another recent study, a extracytoplasmic sigma factor (*σ*X) was shown to remodel the cell wall components of *B. subtilis* in a way that confers phage tolerance in a sporulation-independent manner [57]. Specifically, a stress-response RNA polymerase (*σ*X) activates enzymes that remodel the cell wall including receptors (i.e., techoic acids) that are involved in phage adsorption and plaque kinetics. As a result, a transient phage tolerance response restricts the lysis-induced proliferation of viruses as transcription factors released by infected neighboring cells contact uninfected bacteria. Thus, danger signals can lead to similar outcomes of phage defense in spatially structured population, but for different reasons. Finally, the release and uptake of small peptides can also alter viral-associated cell fate in *B. subtilis*. For example, when infected by SPbeta phages, there is a shift from lysis to lysogeny as the concentration of the communication peptide arbitrium increases during a population-wide infection [58–60]. Future work that explores the resource and signaling landscape modulated by lysis and the relative concentration of infection within spores (i.e., virospores) would provide further insights into the causes and consequences of sporulation-virus feedback in spatially structured habitats across multiple scales.

Irrespective of the molecular mechanisms, sporulation is a complex and energetically costly bacterial strategy that can help cells survive through extreme, often fluctuating, environmental conditions. As such, limitation of viral dispersion by sporulating cells has key implications for the eco-evolutionary dynamics of phage and bacteria. For instance, early sporulation reduces the number of productive infections and the potential for the generation of phage progeny, including host-range mutants. The rapid dispersal and spatial localization of lysis-associated signals could limit phage spread, even in the absence of the evolution of phage-resistant bacterial mutants [22]. However, sporulation may not necessarily constitute a long-term protection against phage infection. For at least some phages that infect *Bacillus*, the lytic reproductive cycle is arrested when a host has already begun to initiate the process of sporulation. Proteins involved in spore development (Spo0A) can bind to regions of the phage genome (0A boxes) that halt the infection cycle. Subsequently, other genes acquired by phages (*parS*) assist with the translocation of the virus genome into the developing forespore, where it resides in an unincorporated state until host germination, at which time the virus can resume its lytic infection cycle [61–63].

In closing, our findings reveal a novel mechanism by which phage infections can be self-limiting leading to a collective defense mechanism that restricts phage spatial spread even when nearby hosts remain available, albeit inaccessible. These findings provide additional context for evaluating the ecoevolutionary dynamics of sporulation in environmental and therapeutic contexts given the link between cellular and population fates. In doing so, it will be essential to consider the consequences of limitations of viral infections across scales including the possibility that collective defense in the near-term may lead to vulnerabilities in the long-term as spores and their viruses re-encounter favorable conditions.

## VI. ACKNOWLEDGEMENTS

We acknowledge Zhongqing Ren, Caroline Dunn, along with the Wang and Kearns laboratories at Indiana University for assistance with microscopy and imaging. We would also like to thank Emily Long for assistance with preliminary data, Thomas Day for advice with image analysis and Adriana Lucia-Sanz for the code review. This work was supported by National Science Foundation (DEB - 1934554 to JTL and DS, DBI - 2022049 to JTL and DEB - 1934586 to JSW, Simons Foundation (930382 to JSW), US Army Research Office Grant (W911NF-14-3071-0411 and W911NF-22-1-0014 to JTL), the National Aeronautics and Space Administration (80NSSC20K0618 to JTL) and the European Research Council (Horizon 2020 research and innovation program - grant agreement No. 740704 to Anastasios M). JSW was supported, in part, by the Chaires Blaise Pascal program of the Île-de-France region.

