## Supplementary Information for "Phage infection fronts trigger early sporulation and collective defense in bacterial populations"

### Supplementary Figures

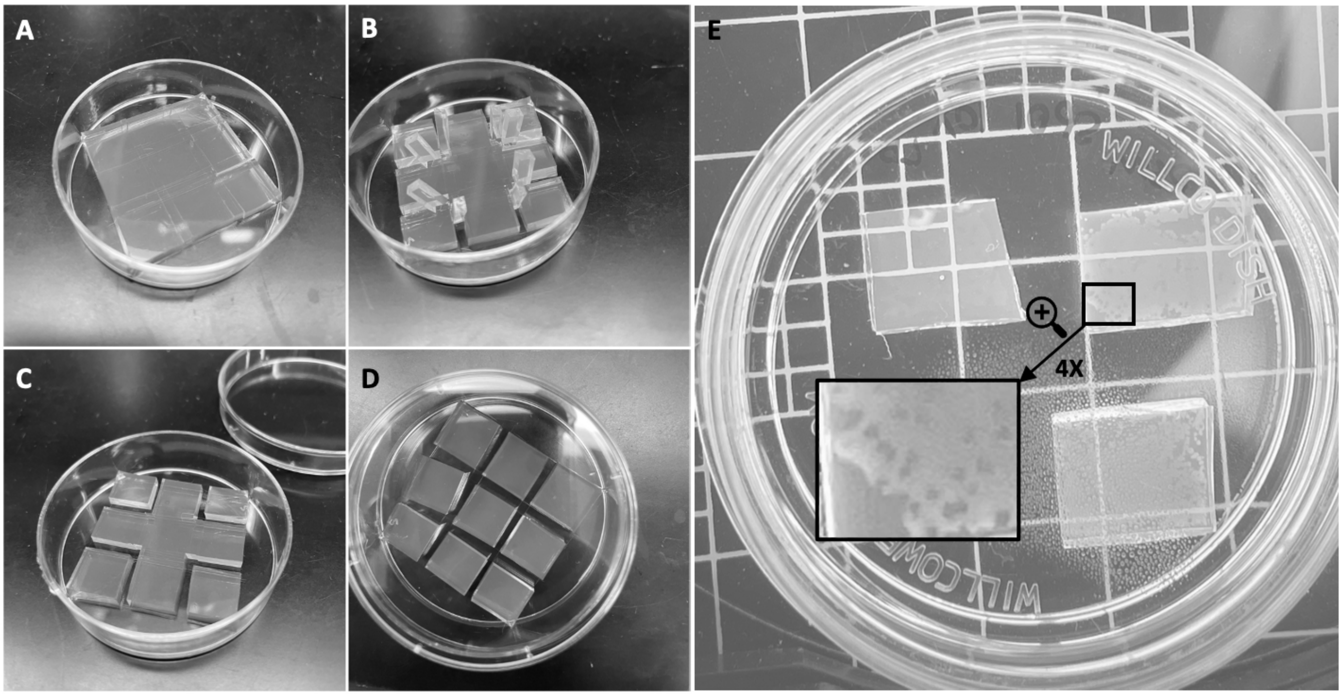


Fig. S1: **Micro plaque assay protocol.** Panels A through D show the preparation of the agar pad for the micro plaque assay in chronological order. The agar layer is cut with a sterile scalpel and the excess agar is removed without moving the square pads. (E) A top-down image of a micro plaque assay is shown. The area enlarged 4 times shows the plaques (darker circular areas) of phage SPO1 on wild type host and DSM media.

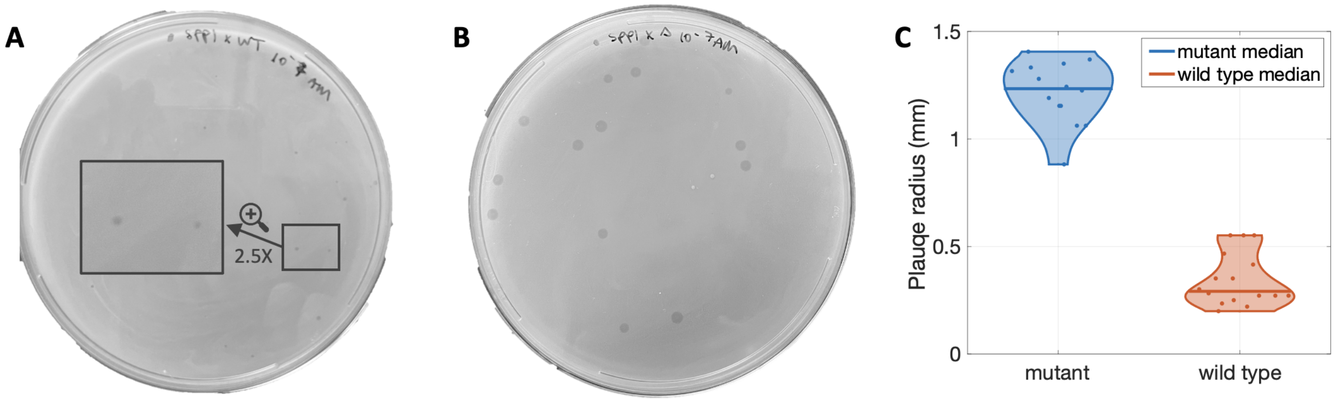


Fig. S2: **Plaque sizes of phage SPP1 with wild type and mutant host.** (A-B) Two entire plaque assays are shown on DSM media. Wild type host is shown in panel A and mutant host in panel B. A small area in panel A is enlarged and shown at 2.5x magnification so that the plaques become visible. (C) Final plaque sizes of the two plaque assays shown in panels A and B. The images were analyzed by hand using FIJI. The plaque sizes (14 data points for mutant and 16 for wild type) were plotted using a violin plot and individual points are shown as a scatter plot. The median mutant and wild type plaque sizes are shown through the horizontal bold lines. Mutant plaques have an average radius of $\approx 1.22mm$ and standard deviation of $0.15mm$ and wild type plaques have an average radius of $0.35mm$ and standard deviation of $0.12mm$. A two sample t-test was performed to reject the hypothesis that the two distributions have equal means with a p-value less than 1e-3.

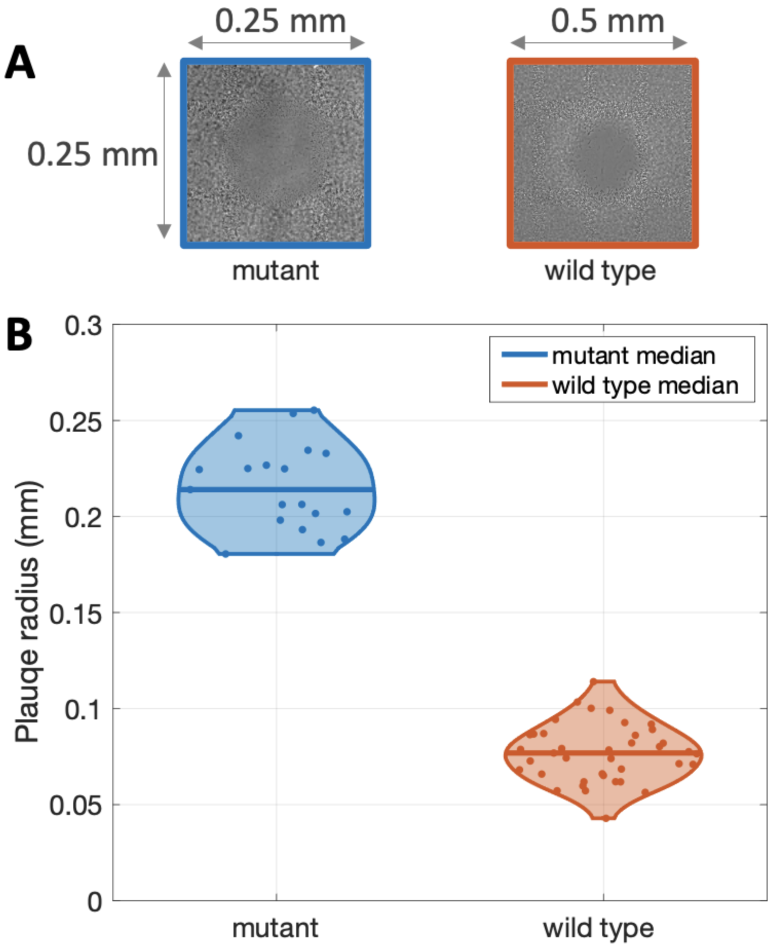


Fig. S3: **Micro plaque sizes of phage SPO1 with mutant and wild type host.** (A) Two micro plaques are shown, both obtained from an inverted microscope (mutant host on the left is taken at 20x magnification and wild type host on the right is taken at 40x magnification). (B) Multiple plaques are measured and the final size distribution is shown for wild type and mutant host (19 data points for mutant and 39 for wild type). The images were analyzed by hand using Fiji. The median mutant and wild type plaque sizes are shown through the horizontal bold lines. Wild type micro plaques have an average radius of 77 microns and 15 microns standard deviation and mutant micro plaques have an average radius of 216 microns and 22 standard deviation. A two sample t-test was performed to reject the hypothesis that the two distributions have equal means with a p-value less than 1e-3.

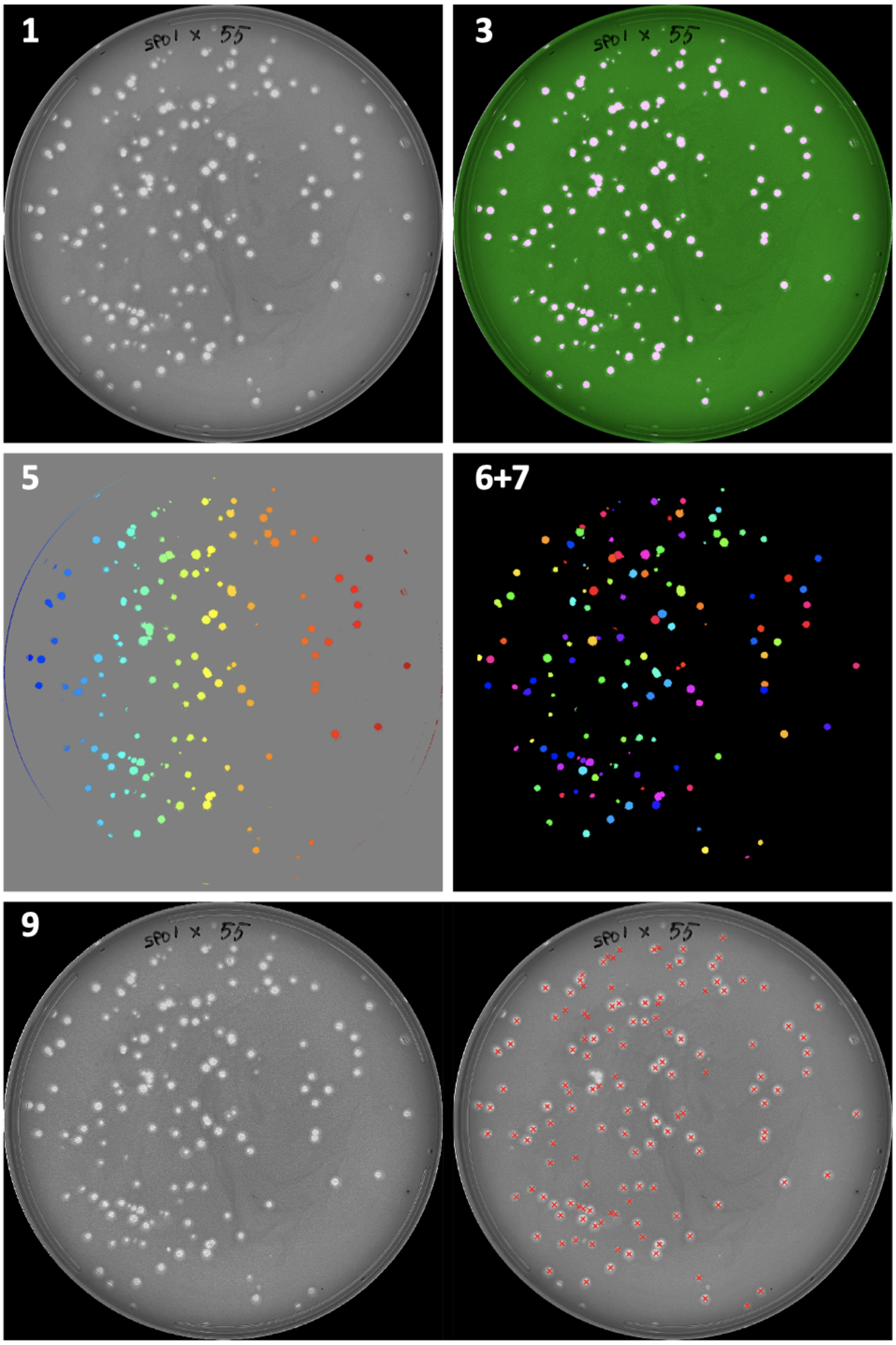


Fig. S4: **Intermediate steps for the final point image analysis algorithm.** Intermediate images for the steps outlined in section VI B 1 are shown. The steps can be described as 1 - original image, 3 - mask from threshold binarization, 5 - watershed algorithm, 6+7-exclusion of false positives, 9-comparison of original image to final plaque centers.

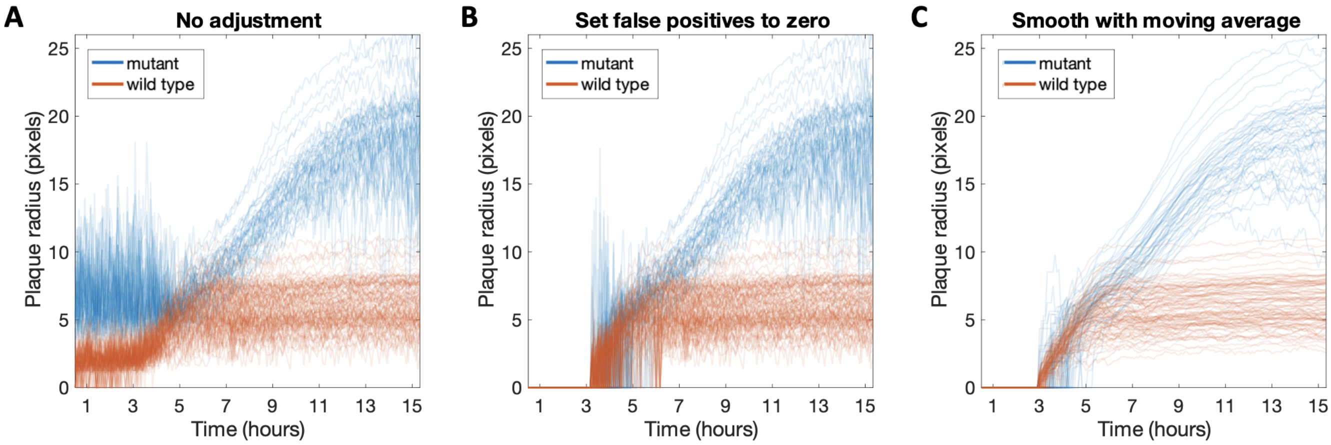


Fig. S5: **Individual plaque sizes at different stages during image analysis** Mutant plaques are shown in blue and wild type plaques are shown in orange across all panels. (A) Raw plaque sizes data are shown after the first part in step 4 in section VI B 2. The plaque size is determined based on a re-scaling and threshold algorithm, hence many early points are false positives in that a plaque is detected when there is none in reality. (B) Plaque sizes at the end of step 4, after setting false positive plaques to zero. All data in the first 33 frames is assumed to be zero. The false positives in the frames 34-70 are determined based on the number of connected components. (C) Final individual trajectories for plaque sizes. The individual growth curves in panel B are smoothed out with a moving average of size 7. These are the trajectories used to determine the mean and standard deviation in figure 2.

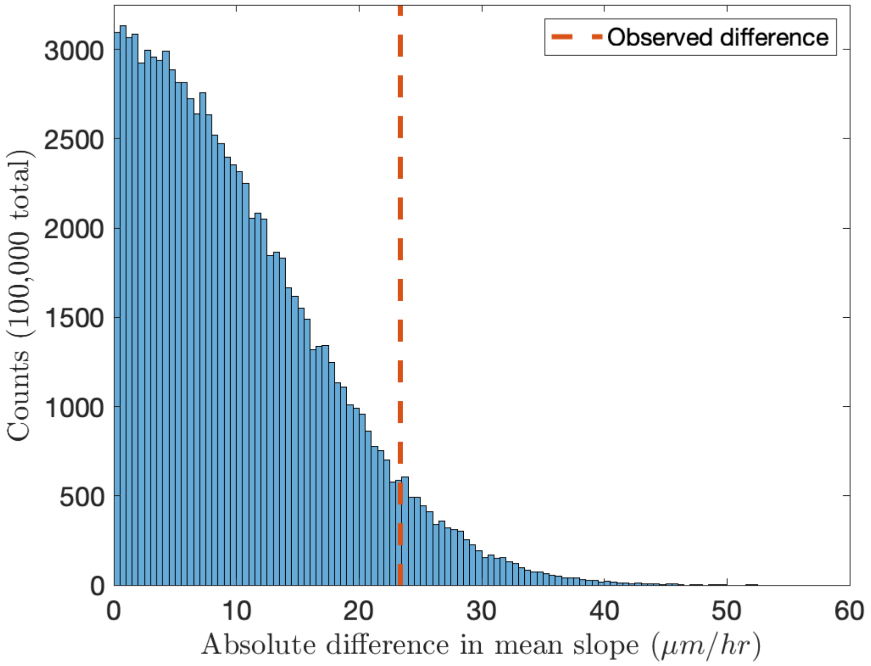


Fig. S6: **Bootstrap analysis of plaque growth rates** Individual plaque growth rates were estimated based on the trajectories from Fig. S5 (details in section VI B 3). Based on the 146 growth rates for wild type and mutant plaques, a bootstrap sampling without replacement was done to show the absolute difference in mean slope. The histogram of the values is shown in blue and the observed difference between the wild type and mutant groups is shown in the dotted orange line (6.6% of bootstrapped differences exceed the observed difference).

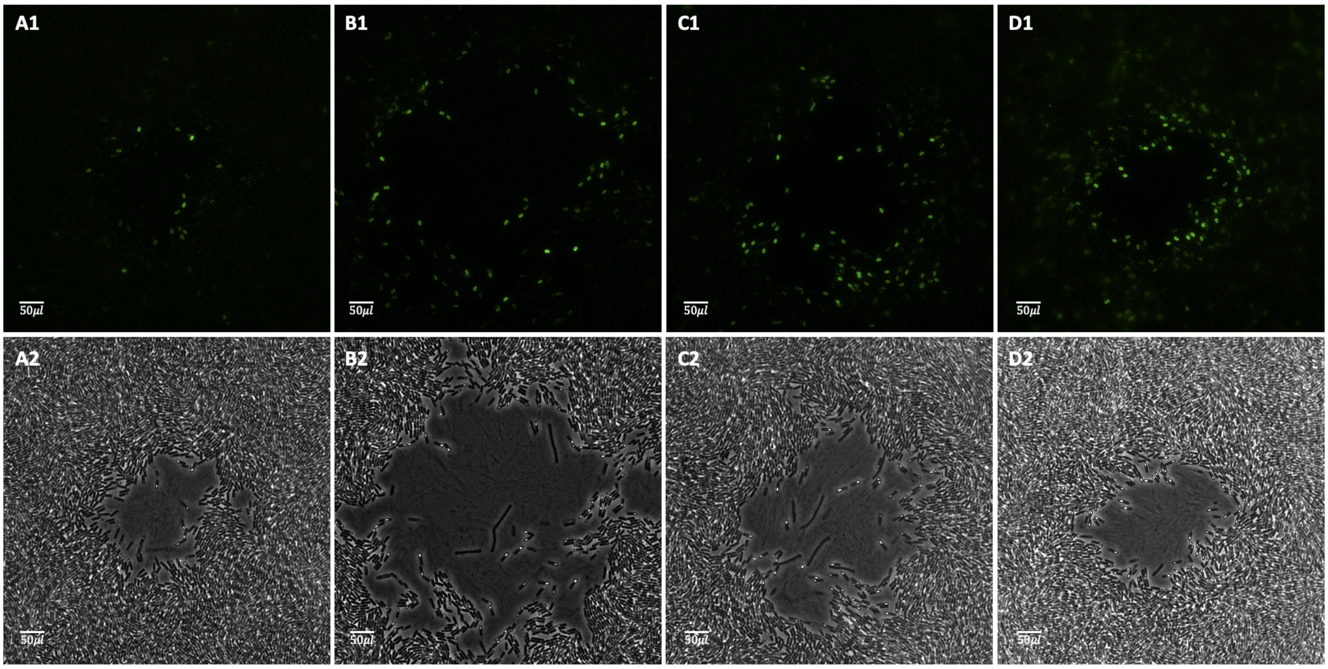


Fig. S7: **Additional images of micro plaque assays at 100x magnification.** The top panels A1-D1 represent the raw GFP image and the bottom ones A2-D2 the raw bright-field corresponding image. All plaques are of phage SPO1 and DSM media. The scale bar in each panel represents 50$\mu m$.

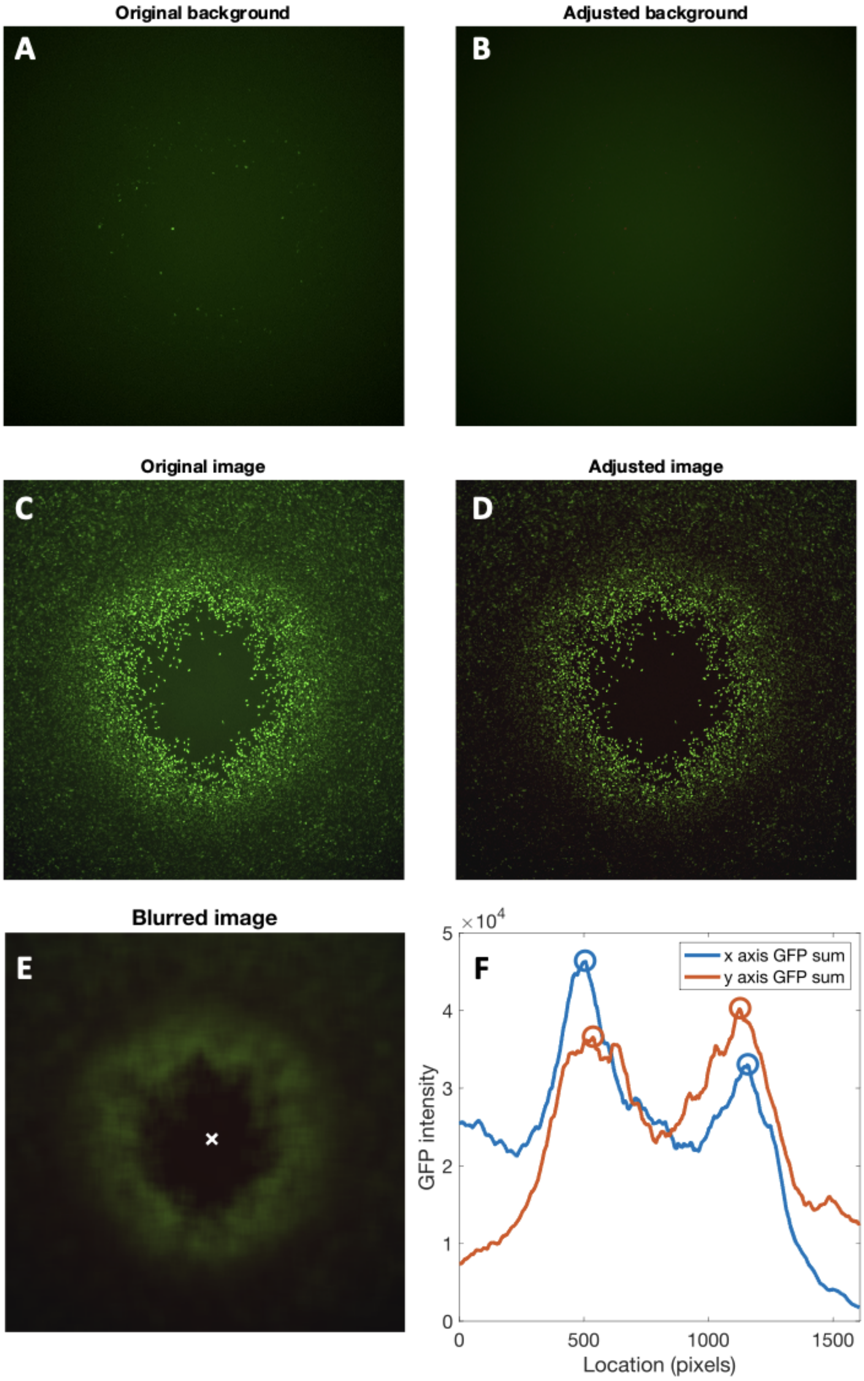


Fig. S8: **Intermediate steps for GFP image analysis** (A) Original image used to calculate the background fluorescence. This image, alongside images in panels B through E, has 1608x1608 pixels. (B) Adjusted background fluorescence in which the bright spots have been replaced with an average of the surrounding area. (C) Original GFP image of the micro plaque. (D) Adjusted GFP image. This image is equivalent to subtracting panel B from C. (E) Adjusted image obtained from applying an averaging filter of size 64x64 to the image in panel D. The white 'x' represents the center of the plaque as computed in panel F. (F) The two solid lines represent the sum and row column of the GFP intensity in panel E. The two peaks for each sum are circled with their respective colors. The midpoints of the blue and orange circles are 831.5 and 832 and represent the coordinates for the plaque center shown in panel E.

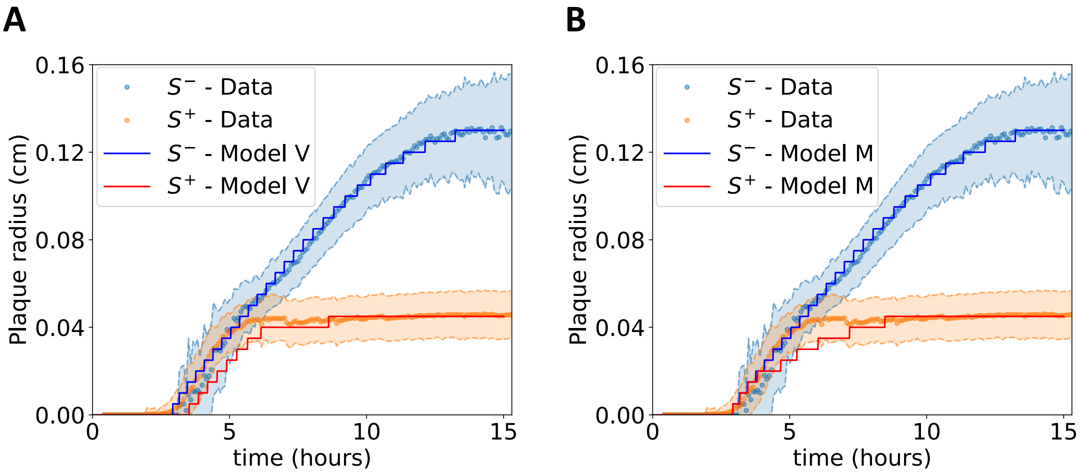


Fig. S9: **Impact of sporulation on plaque size for Models V, and M.** (A-B) An overlay of experimental data and computational simulations showing the time evolution of the mean plaque radius for the $S^-$ and $S^+$ strains in Models V and M, respectively. The results displayed here are for $n_E = 0$ and are consistent with findings for $n_E > 0$ as shown in Fig. 5C,D.

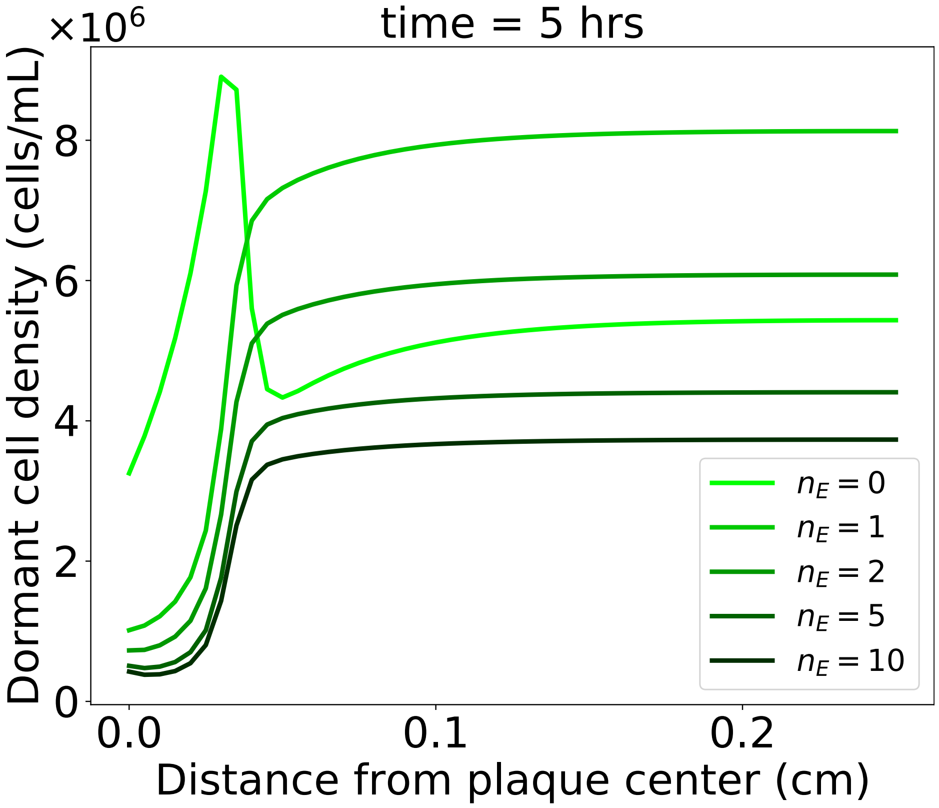


Fig. S10: **The impact of time-delayed transition to dormancy in Model V.** The figure shows how changes in $n_E$ affect the radial profile of dormant cells' distribution for the $S^+$ strain at t = 5 hrs. Darker shades of green represent higher numbers of $n_E$ states. The two extreme cases $n_E = 0$ and $n_E = 10$ correspond to the two cases of Model V presented in Fig. 6D, F.

### Supplementary Text

#### A. GFP-labeled bacterial strain construction

To generate strains producing GFP-labeled spores, we employed DK4422 as the donor strain. DK4422 features a green fluorescent protein fused to a spore coat protein (cotZ), driven by its native promoter, and a red fluorescent protein consecutively expressed by the major sigma factor promoter. These fluorescent reporter genes are integrated into non-essential genes unrelated to sporulation, each accompanied by an antibiotic resistance marker: gfp-cotZ in amyE with chloramphenicol resistance and rfp in ycgO (also known as putP) with MLS resistance.

Due to the chloramphenicol resistance of Δ6 strains, we couldn't use it for selecting the gfp reporter, which is coupled to a Cm resistance marker. However, the proximity of the amyE and ycgO loci in the genome (approximately 20 kB apart) results in genetic linkage between the two engineered reporters. To screen for the cotransformation of both loci, we employed MLS selection for the rfp-mls locus and screened for cotransformation of the gfp locus. To assess cotransformation, we utilized the loss of amylase activity encoded by the amyE gene, which is active in the Δ6 strain but is inactivated by the insertion of the gfp-cotZ gene.

We transformed both Δ6 strains (wild type and mutant) with genomic DNA extracted from DK4422, selecting for growth on MLS. Subsequently, we isolated individual colonies and screened for the loss of amylase activity using the starch-iodine reaction [64]. The presence of GFP in amylase-negative colonies was confirmed through fluorescence microscopy of clones that underwent sporulation in DSM.

To ensure that the transformation and selection processes did not introduce mutations affecting sporulation, we sequenced the genomes of these clones. Library construction (2 x 150 bp), sequencing (Illumina NextSeq500), and sequence analysis were conducted at the Center for Genomics and Bioinformatics at Indiana University. The sequencing reads were mapped to the genomes of the Δ6 strains using breseq. No mutations related to sporulation were identified in either strain in the amyE and ycgO loci. However, in the mutant strain, two mutations, unrelated to sporulation were detected: a nonsynonymous mutation in the aroH gene (V112A), which is involved in the biosynthesis of aromatic amino acids, and a 9bp deletion in the gudB gene, making this cryptic gene active and enabling the utilization of glutamate as a sole carbon source [65].

| Strain | Characteristics | Source (ID) | Ref. |
|---|---|---|---|
| *B. subtilis* 168 Δ 6 | No prophages, $Cm^R$ | BGSC (1A1299) | [66] |
| *B. subtilis* 168 Δ 6 Δ spoIIE | $Spo^-$, No prophages, $Cm^R$ | | [22] |
| DK4422 | *gfp-cotZ* donor to Δ 6 strains<br>amyE::pcotYZ-GFP cat<br>ycgO::PsigA-mRFP mars mls | DK4422 | Kearns Lab |
| DK3895 | $gfp - cotZ$ donor to DK4422 | TMB2112<br>Gifted from lab of prof. Thorsten Mascher at TU-Dresden, Germany | Kearns Lab |
| SPO1 | Lytic phage | ATCC (27370-B1) | [67] |
| SPP1 | Lytic phage | Kearns Lab | |

TABLE I: Table of bacterial and viral strains

#### B. Image analysis

##### 1. Final point analysis

We used a binarization method to extract the plaque sizes from an image of a petri dish. Specific threshold parameters vary between images, however the overall structure of the code is the same. All the analysis was done in MATLAB and a step-wise overview of the code is shown below:

1. Read image and convert to black and white
2. Apply a Gaussian filter to reduce noise
3. Apply a general threshold to obtain a mask
4. Obtain the region properties of the mask
5. Use the build-in watershed algorithm to break apart overlapping plaques
6. Select regions based on circularity and area
7. Discard any additional regions that are not plaques by hand

8. Obtain the areas and centers of the selected regions
9. Perform a final check of the final mask and overlay the centers with the original image

Note that the processed image and binarization was checked against the original image at every time step. We performed steps 3-5 such that no true plaques were being discarded. We used steps 6-7 to ensure that any false positives are not counted. Intermediate images for some of the steps are shown in Figure S4.

##### *2. Time lapse analysis*

To analyze the time lapse of plaque growth, we first analyzez the last frame to identify the plaques. We then used the plaque centers and axes to track the size of the plaques in previous frames. This process can be divided into 5 steps described in detail below.

**Step 1. Identify plaque centers in last frame**

A circular section was cropped from the original video using code adapted from MATLAB Central Answers to obtain a time lapse for each petri over time (link : https://www.mathworks.com/matlabcentral/answers/567441-how-to-crop-circle-from-an-image). We then used the same protocol as the one described in section VI B 1 to identify the centers and axes of the plaques in the last frame.

**Step 2. Compute camera movement**

To obtain a video of the plaques over time, we set a camera to capture a top-down image of the petri dishes every 5 minutes. We note that the camera was not perfectly still and that there was random movement in the x and y direction between frames. We corrected for this noise by tracking the edges of the white LED screen. The full image was cropped to contain just the corner of the LED screen. The cropped image was then binarized and 10 rows and columns were used to find the transition between the white screen and the background. The first black pixel for each row marked the transition and was used to adjust for displacement in the x direction. Similarly, the first black pixel for each column marked the transition and was used to adjust for displacement in the y direction. In the end we obtained 10 points marking a vertical edge and 10 points marking a horizontal edge for each frame.

**Step 3. Identify search window based on centers, camera movement and axes lengths**

Instead of analyzing the entire petri dish at every time step, we used the information about the plaque location and size to analyze a narrow window for each plaque. For each frame we took the mean of the 10 points marking the vertical edge and 10 points marking the horizontal edge. The mean values for the two edges were compared to the ones in the last frame to obtain the camera movement in the $x$ and $y$ direction with respect to the last frame. The centers of the plaques were adjusted to account for this camera movement. Thus, starting from the center of the plaques in the last frame, we obtained the center of the plaques in all the previous frames.

Lastly, for each frame and each plaque we defined a rectangular area centered at the adjusted plaque center. This rectangle has the width and length equal to the two major axes of the final plaque size. Since plaques only grow over time, it means that the plaque at the current frame is contained within the rectangle.

**Step 4. Calculate plaque in each rectangle for all plaques and frames**

Having defined a search area for each plaque and each frame, we were left to differentiate between plaque and lawn in each rectangle. Since the contrast between plaque and lawn is not high enough in earlier time points, we could not use a general threshold for this algorithm. Instead we wrote a short script which is used for both wild type and mutant time lapse. Given rectangle $X$, we made the following operations:

$X = \text{imadjust}(X)$ ▷ rescale of intensity values
$X_{BW} = X > 160$ ▷ threshold binarization
$X_{BW} = \text{fillholes}(X_{BW})$

The plaque sizes after this analysis are shown in Figure S5 panel A, however we note that there are plenty of false positives, i.e. areas that are not plaques but are counted as such. To fix this issue, we divided the 179 frames in 3 time intervals and adjusted the false positives to zero based on the number of connected components $n_{CC}$ in $X_{BW}$. A large number of connected components is an indication of a false positive. We selected the $n_{CC}$ threshold by plotting the number of connected components for each plaque over time. A brief version of the code is shown below:

**if** $i < 34$ **then**
  plaque size = 0 ▷ Plaques are not visible yet
**end if**
**if** $34 \leq i < 70$ **then**

    **if** $n_{CC}$ > threshold **then**
        plaque size = 0 ▷ Set false positive to zero
    **else**
        plaque size = largest connected component in $X_{BW}$
    **end if**
**else**
    plaque size = largest connected component in $X_{BW}$
**end if**

The plaque sizes after adjustment for false positives can be seen in Figure S5 panel B.

**Step 5. Raw data and smoothing function for each plaque**
We expect plaques to grow monotonically over time, however there is noise in the original video and in our image analysis algorithm. To reduce the noise and emphasize the overall trend of the plaque measurements, we took a moving average of size 7 for each plaque trajectory. These are the final plaque growth trajectories we use for the main text and they are also shown in Fig. S5 panel C.

##### *3. Bootstrap analysis*

Using the trajectories in Fig. S5 panel C, we found the individual plaque growth rates by fitting a linear model to the plaque sizes during enlargement phase. For each plaque, we defined the enlargement phase based on the size of the plaque at the end of the time-lapse experiment. Because wild type plaques are smaller than mutant plaques, we used a wider window for the wild type strain as follows: 30-70% for wild type plaques and 50-70% for mutant plaques. We then fit a linear model to each individual growth curve and obtained a total of 146 linear growth rates (46 for wild type and 100 for mutant). To test whether the growth rates for wild type and mutant are statistically significant, we performed $10^5$ bootstrap sampling steps without replacement to divide the 146 growth rates into two sets of size 100 and 46. For each bootstrapping step we computed the absolute mean difference between the two groups and showed the results against the original difference in Fig. S6. We obtained that the plaque growth rates of wild type and mutant are different with a p-value of 0.066.

##### *4. GFP analysis*

We used a set of imagining analysis techniques to obtain the GFP intensity based on the distance to the center of the plaque shown in figure 3. The process can be described in three steps outlined below:

**Step 1. Calculate background fluorescence and subtract it from original image**
The original GFP image (Figure S8 panel C) has a background fluorescence level that is higher at the center of the image. To account for this, we used another GFP image with minimal fluorescent cells shown in panel A as a measure for background GFP intensity. We took out the bright spots in the original background and filled them in to obtain an adjusted background fluorescence shown in panel B. Finally, subtracting the adjusted background from the original image yielded the adjusted GFP image in panel D which was used for analysis in the following steps.

**Step 2. Find center of the plaque**
Once we obtained the adjusted GFP image, we used a square averaging filter of size 64x64 to obtain an adjusted image shown in Figure S8 panel E. Using the adjusted image, we computed the GFP sum for each row and column and noticed that there were two peaks for both the column and row sums shown in panel F. These peaks correspond to the edges of the plaque and we selected the midpoint for the peaks to represent the x and y coordinate for the center of the plaque.

**Step 3. Plot GFP intensity based on distance to center**
Once we found the center of the plaque, we computed the distance from each other pixel ($1608^2$ pixels in total) to the center. The pixels were grouped in 400 bins based on the calculated distance and a mean and standard deviation was obtained for each bin. The mean and standard deviation are the final measurements shown in figure 3 panel C.

#### C. Mathematical modeling and simulations methods

##### 1. Numerical simulations

We simulate the PDEs through a finite difference method implementation on a square lattice. Each lattice sites consists of a square with $\Delta x = 50\mu$m$= \Delta y$ with 100 sites per side to represent a total length $L = 0.5$ cm. In this implementation, for each time step, the contribution to the derivatives given by the local reaction terms are computed in each site, based on the state values in that site at that given time, as when integrating ODEs. The spatial derivatives $\frac{\partial}{\partial x}, \frac{\partial}{\partial y}$ present in the diffusion terms, which couple different positions, are discretized on the square lattice. We use the following stencil matrix to discretize the Laplacian operator ($\nabla^2 = \frac{\partial}{\partial x}^2 + \frac{\partial}{\partial y}^2$) in the diffusion terms:

$$\text{Kernel} = \begin{pmatrix} 0.25 & 0.5 & 0.25 \\ 0.5 & -3 & 0.5 \\ 0.25 & 0.5 & 0.25 \end{pmatrix}$$

This stencil needs to be convoluted with the state field. For instance the diffusion term of resources $D_R \nabla^2 R$ at a site (x,y) can be obtained by discretizing spatial derivatives to first order with respect to first- and second-neighbor sites:

$$\begin{aligned} D_R \nabla^2 R(t,x,y) &= \frac{D_R}{\Delta x^2} \begin{pmatrix} 0.25 & 0.5 & 0.25 \\ 0.5 & -3 & 0.5 \\ 0.25 & 0.5 & 0.25 \end{pmatrix} \cdot \begin{pmatrix} R(t,x-\Delta x,y+\Delta y) & R(t,x,y+\Delta y) & R(t,x+\Delta x,y+\Delta y)) \\ R(t,x-\Delta x,y) & R(t,x,y) & R(t,x+\Delta x,y) \\ R(t,x-\Delta x,y-\Delta y) & R(t,x,y-\Delta y) & R(t,x+\Delta x,y-\Delta y) \end{pmatrix} \\ &= \frac{D_R}{\Delta x^2} \Big( -3R(t,x,y) + 0.5\Big(R(t,x+\Delta x,y) + R(t,x-\Delta x,y) + R(t,x,y+\Delta y) + R(t,x,y-\Delta y)\Big) + \\ &+ 0.25\Big(R(t,x+\Delta x,y+\Delta y) + R(t,x-\Delta x,y+\Delta y) + R(t,x+\Delta x,y-\Delta y + R(t,x-\Delta x,y-\Delta y)\Big)\Big) \end{aligned}$$

Diffusion of viruses and other molecules is given by the same equation by accounting for their specific diffusion constant. Together with the site-specific reaction terms, this step completes the computation of time derivatives given all the states variables at each time point. We can then update the system for 15 hrs using time steps of 0.01 hrs, by solving an explicit Runge-Kutta method of order 5(4) (Scipy's default initial value problem solver for a system of ordinary differential equations) [68, 69].

##### 2. Parameter estimations

- Host-growth

  A maximum growth rate of $r_{max} = 2/hrs$ corresponds to a doubling time of approximately 20 min [70]. We constrained a set of parameters to a plausible order of magnitude based on back-of-the-envelope calculations and comparisons to the elements from the literature. We set the initial amount of resources (in rescaled units) as $\tilde{R_0} = 10^9$ cells/mL, which sets the carrying capacity for bacteria growth. Several parameters were obtained *de novo* in this work, either based on the from the experimental protocols, or inferred (qualitatively) to reproduce the experimental results. The resource to bacteria conversion rate order of magnitude $\epsilon$ is estimated by noting that a cell contains typically 200 fg of Carbon [50] and assuming that half of nutrient weight is due to Carbon, that means it takes about 400 fg of processed nutrients to make up a cell. Assuming a cell metabolizes to growth a bit less than half of the uptaken nutrient, we get $\epsilon \approx 10^{-6} \mu g/cell$. The Monod constant $K_g$ corresponds to the resource level where the growth rate is half of its maximum value. To reproduce the experimental results we select a Monod constant that is approximately three times larger than the initial resources, with the growth rate changing almost linearly with resources. Since the maximum plaque radius is under 0.2 cm, the size of the lattice is set to $L = 0.5cm$. We estimate the initial number of cells per plate to be $100\mu l$ of culture at OD 0.5 $\approx 2 \times 10^8$ cells/mL, corresponding to $\approx 2 \times 10^7$ cells. Given a bacterial lawn height of 400 $\mu m$ and plate area of $\pi \cdot 4.5^2 cm^2$, we estimate the volume of a plate where bacteria grows to be about $\pi \cdot 4.5^2 \cdot 0.04 cm^3 \approx 2.5ml$. We thus obtain an estimate for $S_0 \approx 2 \times 10^7 cells/2.5ml \approx 10^7 cells/ml$.

| Parameters common to models R, V and M | | | | |
|---|---|---|---|---|
| *Variable* | *Meaning* | *Value* | *Unit* | *Source* |
| $R_0$ | Initial resource level | $10^3$ | $\mu$g/mL | This work |
| $\tilde{R}_0$ | Rescaled initial resource level | $10^9$ | (cells)/mL | This work |
| $S_0$ | Initial cell density | $10^7$ | (cells)/mL | This work |
| $V_0$ | Initial viral density | $10^6$ | (viruses)/mL | This work |
| $r_{max}$ | Maximum bacterial growth rate | 2 | hrs$^{-1}$ | [70] |
| $\varepsilon$ | Resource to bacteria conversion rate | $10^{-6}$ | $\mu$g/(cell) | This work |
| $K_g$ | Monod constant | $3.2 \cdot R_0$ | $\mu$g/mL | This work |
| $\tilde{K}_g$ | Rescaled Monod constant | $3.2 \cdot \tilde{R}_0$ | (cells)/mL | This work |
| $\phi$ | Infection rate | $1.8 \cdot 10^{-8}$ | mL/(hrs $\cdot$ (virus)) | [22] |
| $\eta_{max}$ | Maximum latent rate | 4/3 | $hrs^{-1}$ | [71] |
| $d_{max}$ | Maximum transition to dormancy rate | 1 | $hrs^{-1}$ | This work |
| $\beta$ | Burst size | 100 | (viruses/cell) | [72] |
| $\omega$ | Viral decay | 0.001 | hrs$^{-1}$ | [72] |
| $D_R$ | Resource diffusion constant | $4 \cdot 10^5$ | $\mu$m$^2$/hrs | [41] |
| $D_V$ | Phage diffusion constant | $2 \cdot 10^3$ | $\mu$m$^2$/hrs | This work |
| $n_I$ | Number of infected states | 10 | | [41] |
| $s$ | "Sharpness" of transition to dormancy | $10^{-7}$ | mL/(cells) | This work |

TABLE II: **Parameters common to models R, V, and M.** This table lists the variables used across models R, V, and M. Each row corresponds to a specific parameter while columns present the respective information including the parameter's symbol, meaning, value, unit, and source. Additional parameters unique to models V and M, which build upon Model R, are discussed exclusively in the Mathematical Models and Simulation section and in the Parameter estimations subsection.

- Viral infection

  Since in our deterministic models a plaque starts from exactly one virus (compatible with the standard definition of PFU) and the volume of one square in the lattice is $Vol = 50\mu m \cdot 50\mu m \cdot 400\mu m = 1000000\mu m^3 = 10^{-6}ml$, we get that $V_0 = 1/Vol = 10^6 ml^{-1}$. The max latent rate $\eta_{max}$ is given by [71], while for the burst size $\beta$ [72], the viral decay $\omega$ [72] and the resource diffusion constant $D_R$ [41] we selected plausible values that are commonly used in similar phage-bacteria ecology studies. The infection constant $\phi$ for the Bacillus Subtilis-SPO1 pair is close to but smaller (about an order of magnitude) than the value reported by [22]. The values of $d_{max} = 1/hour$, $n_I, n_E$ when $> 0$, and the absolute magnitude of $\mu$ and $m$ were set to reproduce the experimental dynamics semi-quantitatively.

- Dormancy

  For the transition to dormancy we selected values for $\lambda$ that correspond to a process taking approximately 45 minutes from a signaling event, which is close to the reported time for commitment to sporulation [73]. Since dormancy is the "last resort" for a bacterium under stress, we select the threshold to dormancy $\sigma$ to be a small percentage of the Monod constant, at about the point where cells grow 3 to 5 times slower than normal. In order to mimic a precise signaling process that triggers sporulation at critical resource level, we set the shape parameter of the resource-triggered dormancy Hill function, $s$, to be two orders of magnitude smaller than the carrying capacity $\tilde{R}_0$. Finally we assume that the product between $\mu$ and $m$ is similar to that of the infection rate $\phi$ and the burst size $\beta$, that is, the size of a peptide chain will correlate with both the ability of a cell to synthesize it in large numbers and with the rate at which a cell will bind to it and adsorb it, since in a diffusion limited reaction the adsorption rate will scale linearly with the diffusion constant [74] which is inversely proportional to the particle size due to the Einstein-Stokes equation.

- Spatial diffusion

  To estimate the phage diffusion constant $D_V$ we consider phage to be about two orders of magnitudes larger than the molecular components constituting resources. Using Einstein-Stokes relation yields a value for $D_V$ that is two orders of magnitude smaller than the resource diffusion constant. As a cross check we note that this is compatible with values proposed in literature, albeit for different phages [75]. We selected a molecule diffusion constant $D_M$ to be equal to the resource diffusion constant $D_R$ since we expect the molecule to be of similar size to resources.
